# Hyperacute Response Proteins (HARPs) synthesized on γ-tubulin-FTO-MARK4 translation microdomains upon exposure to stress, regulate stress response in cancer

**DOI:** 10.1101/2025.01.20.633998

**Authors:** Bruno Saleme, Saymon Tejay, Paul Dembele, Rabih Abou Farraj, Yongneng Zhang, Yongsheng Liu, Aristeidis E. Boukouris, Sotirios D. Zervopoulos, Alois Haromy, Yuan-Yuan Zhao, Shelly Braun, William Saleme, Xuejun Sun, Richard Fahlman, Mark Glover, Adam Kinnaird, Gopinath Sutendra, Evangelos D. Michelakis

**Affiliations:** Department of Medicine, University of Alberta, Edmonton Canada; Department of Biochemistry, University of Alberta, Edmonton Canada; Department of Oncology, University of Alberta, Edmonton Canada; Department of Surgery, University of Alberta, Edmonton Canada

## Abstract

Compared to normal, cancer cells are particularly resistant to stress, and their immediate response to stress is critical for their subsequent multilayered adaptation programs which pose a major clinical challenge. With unbiased proteomics and transcriptomics analysis, we identified a list of HARPs synthesized from pre-existing mRNAs within 20 min of diverse stresses in A549 cancer cells, despite the known suppressed global translation in stress. HARP mRNAs were translated on microtubule-associated translation microdomains (MATMs) located on γ-tubulin, that host FTO and specialized cytoskeletal ribosomes, structurally and functionally distinct from ER and cytosolic ribosomes. FTO exited the nucleus immediately after stress and was activated by the microtubule-associated stress kinase MARK4 via T6 phosphorylation. Activated FTO demethylated a translation-inhibiting mRNA methylation (m6A) signature, facilitating compartmentalized HARP translation on MATMs, while non-HARP mRNA remained inhibited. FTO or MARK4 inhibition suppressed HARP synthesis and increased apoptosis post various stresses, including chemotherapy. These data were confirmed in 4 additional cancer cell lines and normal fibroblasts. Using the Protein Atlas database, we found that high levels of our identified HARPs had on average a 35% decrease on patient 5-year survival in prevalent and resistant cancers (breast, lung, liver, pancreas). γ-tubulin, FTO and MARK4 are therapeutic targets for many cancers, through their ability to comprehensively promote HARPs translation, a potential Achille’s heel for cancer’s resistance to physiologic or therapeutic stress, offering a new window in stress biology.

## Introduction

Cancer cells are constantly exposed to a variety of microenvironment stresses (e.g., nutrient deprivation early in carcinogenesis due to suboptimal tumor vascularization) or therapy-induced stresses, (e.g., chemotherapy, radiation therapy), that are sufficient to irreparably damage benign cells, yet cancer continues to be a leading cause of death world-wide^1^. This is due to cancer’s ability to resist stress via numerous mechanisms, including increased expression of drug efflux pumps, enhanced ability to repair DNA and remodel their metabolism and cytoskeleton to exit the cell cycle and develop features of stemness, among others^2, 3^. In addition to enhanced ability to survive the initial stress, cancer cells have a remarkable ability to adapt and reprogram so with each stress cycle, they become more resistant to subsequent stresses. For example, following development of resistance to one chemotherapy, they develop resistance to other chemotherapies they had not been exposed to before^4^. For cancer cells to acutely survive a new stress and adapt, there likely exists a comprehensive stress response program that gets activated early after stress exposure to first ensure survival, then repair the damage and subsequently reprogram toward long-term adaptation.

We hypothesized that immediately upon exposure to stress, cancer cells translate a set of Hyperacute Response Proteins (HARPs) that reprogram them to survive, repair, and adapt to future stresses. To ensure a timely response, HARPS are likely to be synthesized from pre-processed and localized mRNAs, bypassing transcription, mRNA processing, and mRNA transfer to translation sites, which altogether can take hours, independent of gene length which also greatly influences timing^5, 6^. HARP translation should be inhibited at baseline prior to stress to avoid unnecessary stress-reprograming in the absence of stress. Thus, we speculated that the mechanism of HARP translation would involve the methylation of n-6 adenosine (m6A), a common post-transcriptional modification on mRNA, which has been shown to suppress translation rates when localized to the coding regions of mRNA, via dissociation of tRNAs and prevention of elongation, or by decreasing the stability of mRNA.^7–9^. Increasing evidence implicates m6A in stress response^10^, as well as in cancer aggressiveness, resistance to therapy and metastasis^11^. All components of the m6A axis, from writers like the main methyltransferase that methylates mRNA in the nucleus (METTL3)^12^, to erasers like the m6A demethylase *Fat mass and obesity protein* (FTO)^13^, to readers like YTHD1^11^ have been shown to play some role in cancer biology via various mechanisms. The lack of a comprehensive mechanism makes the role of the m6A axis in cancer resistance to stress difficult to understand.

We speculated that HARP mRNAs may be decorated by an m6A signature that **a)** supresses their translation at baseline in the absence of stress and **b)** allows them to be recognized by m6A readers that facilitate their transfer to specialized translation microdomains, thereby facilitating their selective translation in stress. Sub-cellular compartmentalized translation is best suited for the selective HARP mRNA translation, while translation rates are known to be suppressed during acute stress as a means of saving energy, since translation is the cell’s most energy-demanding function. Indeed, subcellular translation microdomains and ribosomal heterogeneity have been shown to play important roles in stress biology and have expanded our understanding of translation diversity^14–19^. We hypothesized that demethylation of a translation inhibitory HAPR-specific m6A signature during hyperacute stress provides a selective translation signal, mediated by cytoplasmic activation of FTO (which is known to shuttle outside the nucleus^20, 21^). These processes likely occur on translation microdomains perhaps hosting specialized ribosomes. The removal of an m6A HARP-specific signature, may allow HARP mRNAs to be translated at higher rates immediately after stress onset, while the non-HARP mRNA translation rates would remain supressed during stress.

We used the widely studied A549 cancer cells, which are known to develop resistance, and exposed them to a pulse of mild UV irradiation (25 J/m^2^), as a model of genotoxic stress, or metabolic stresses induced by 2DG (2-deoxyglucose) which inhibits glucose utilization, or by deprivation of the essential amino acid methionine. All these stresses are relevant to the stress that most solid tumors are exposed to, either at early stages or during cancer treatments^22^. We focused on events occurring within the first 20 minutes of stress (i.e., hyperacute stress), a stage of stress often not specifically studied in stress biology research. These findings were all reproduced in other commonly used cell lines. To identify HARPS, we used unbiased proteomics and transcriptomics, and to visualize HARP mRNAs within translation microdomains, we used RNA hybridization and high-resolution confocal microscopy. We also used imaging (confocal and transmission electron microscopy with phosphorus mapping) along with ribosome sequencing to study ribosomal diversity.

## RESULTS

### Microtubule Associated Translation Microdomains (MATMs) increase acutely after stress

Using confocal microscopy, we found that FTO **(Fig. 1A)** but not METTL3 **(Fig. S1A)**, exits the nucleus and enters the cytoplasm within 20 min of UV stress. We also confirmed this by immunoblots **(Fig. S1B).** To visualize active translation in the cytosol during the first 20 min post stress, we labelled newly translated proteins with O-Propargyl puromycin (OPP^23^), an amino acid analogue that can be incorporated in the nascent peptides as they are synthesized, which we added immediately after UV stress for a duration of 20 min. In the absence of stress, we detected a uniform diffuse pattern, which within 20 min of UV and 2DG stress, decreased significantly and was eliminated by the translation inhibitor cycloheximide (**Fig. 1B**), confirming the global decrease in protein synthesis after stress. The overall decrease in protein synthesis during acute stress was also supported by the decrease in radioactive methionine labelling with S35 resulting in an average reduction of global translation of 45% within 20 min (**Fig. S1C**). While the total OPP diffuse signal decreased, distinct OPP foci appeared within this 20 min period post UV and increased in number, in both UV and 2DG stresses (**Fig. 1C**). These active translation foci were distinct from stress granules, detected with staining for TIAR (a stress granule marker), also known to increase acutely after stress (but do not play a role in active protein synthesis)^24^. As expected, the location of stress granules had no significant correlation with OPP translation foci (**Fig. S1D).** Using high resolution immunofluorescence of FTO, we found that cytosolic FTO was associated with β-tubulin, suggesting that microtubules may play a role in the acute microdomain translation hubs after stress **(Fig. 1D)**. This is in keeping with a wealth of data on the importance of cytoskeleton in regulating protein synthesis and stress signalling^25–27^. We found that OPP foci were associated with β-tubulin specifically at bifurcation points with a characteristic “Y” pattern, suggesting colocalization with γ-tubulin (**Fig. 1E-F)**^28–31^. Immunofluorescence staining of FTO, OPP, S6 (a core ribosomal protein) and β-tubulin confirmed the overlap between these elements at the “Y” junctions of microtubules after stress (**Fig. 1G).** Due to limitation of immunofluorescence resolution (i.e., 180nm on our Airyscan confocal system), we further confirmed the colocalization of FTO, S6, γ-tubulin and OPP, using the proximity ligation assay (PLA, a technique that combines antibody binding specificity with DNA probe amplification, allowing colocalization detection within 30 nm^32^) and found a near perfect colocalization between FTO/γ-tubulin and OPP foci, which increased significantly with UV stress (**Fig. 1H)**. Similar findings to OPP were seen with L-Azidohomoalanine (L-AHA, or AHA) labelling of newly synthesized proteins^33^, which is a methionine analogue that does not normally exist in cells but can be utilized by translation machinery, allowing for the detection of newly synthesized proteins by adding it to culture media immediately after UV stress. Similar to OPP, the L-AHA signal colocalized with the PLA signals of FTO/γ-tubulin **(Fig. S1E)**. We named these microdomains “Microtubule Associated Translation Microdomains (MATMs). Overall, these data show that amidst the decreased translation in the first 20 min of diverse stresses, new protein synthesis takes place within γ-tubulin-linked MATMs. We then studied the potential mechanism by which cytosolic FTO may be activated at stress.

**Figure 1:**
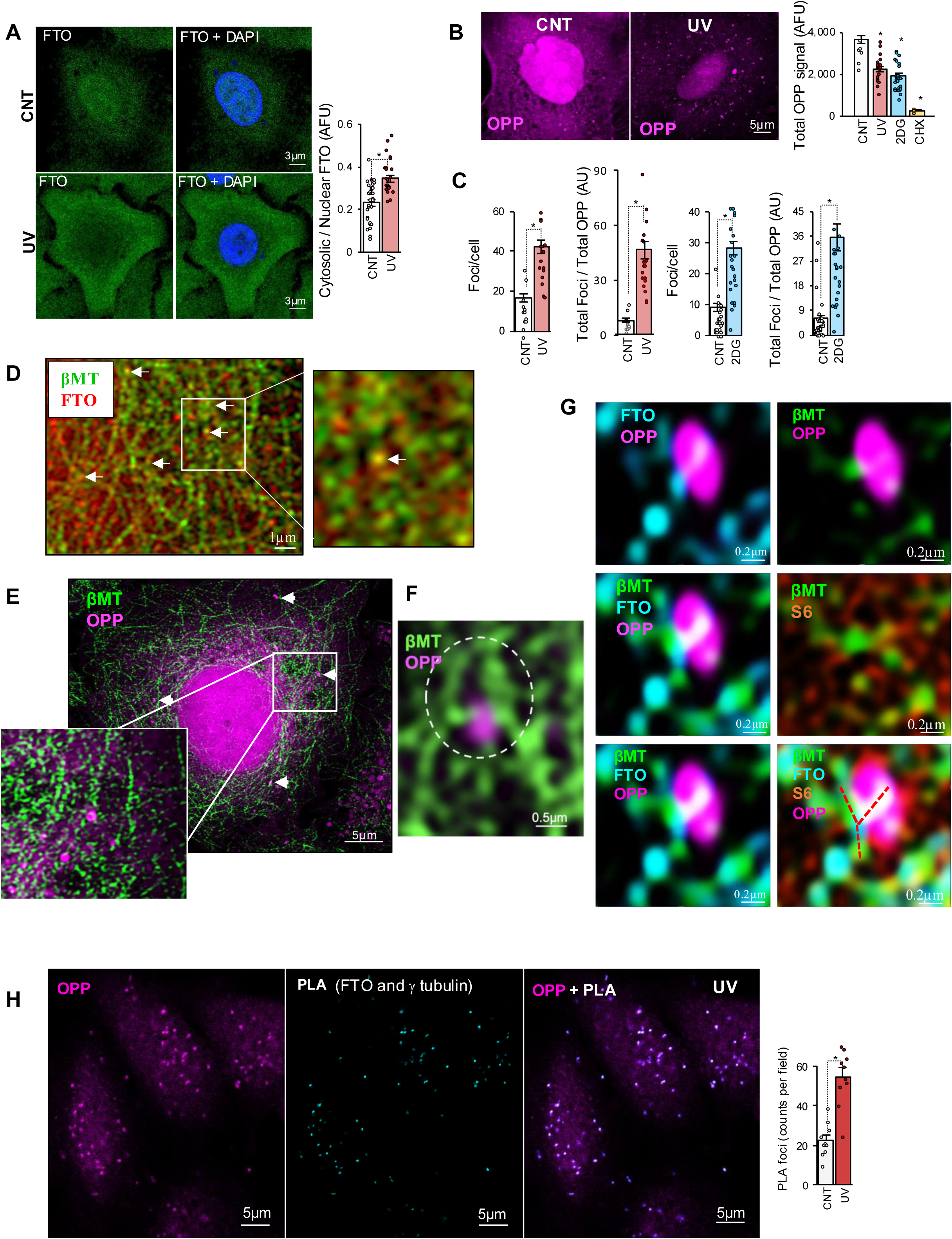
Microtubule Associated Translation Microdomains (MATMs) increase acutely after stress. A) Immunofluorescence staining of FTO (green) and DAPI (blue) in control and UV-treated cells showing FTO signal in the nucleus and cytosol in control (CNT) and UV-treated cells. Mean data and SEM on the right show increased ratio of cytosolic to nuclear FTO levels 20 minutes after UV treatment, suggestive of stress-induced acute FTO nuclear exit. n=3 experiments with data from each plate shown as individual data points; *p<0.00001 to control. B) Staining with OPP shows that while the overall diffuse OPP signal (which signifies active translation), decreases significantly within 20 min of UV stress, distinct well-demarcated foci of OPP signal increase in number during the same time, suggesting an increase in active translation within foci/microdomains. Mean data of total OPP signal is shown on right with decreased signal with UV or 2DG stress and near complete loss of signal with cycloheximide (ribosomal inhibitor) treatment. n=3 experiments with data from each plate shown as individual data points; *p<0.00001 to control. C) Mean data of the ratio of the number of OPP foci normalized to the total cellular OPP signal, or foci per cell, in UV and 2DG-treated cells, compared to control, show increase in foci count in both stresses. n= 5 experiments, p<0.0001. D) Immunofluorescence staining of FTO (red) and β tubulin marking microtubules (MT), green) showing a tubular pattern of MT with FTO overlap. Image on the right shows magnification of inset with overlap shown in yellow as highlighted by white arrow. E) Immunofluorescence staining showing OPP (magenta) and β-microtubules (βMT, green) in UV-treated cell showing the dense translation foci colocalization with microtubular patterns. White arrow heads show the colocalization points. Inset is a magnification of the area enclosed within the white box. F) An example of high magnification of OPP (magenta) βMT (green) showing OPP foci over a β-tubulin branching point. G) Simultaneous immunofluorescence staining with several antibodies (OPP-magenta, βMT-green, FTO-teal, ribosomal protein S6-orange) showing that all these components cluster together. We named these active translation microdomains Microtubule Associated Translation Microdomains (MATMs). The location of OPP on microtubule junctions/bifurcation points is shown by the white co-localization color in the bottom images and highlighted by a red “Y” in the bottom right. H) Immunofluorescence staining of OPP (in magenta) and the PLA signal between FTO and γ-tubulin (in teal) in UV-treated cells showing strong colocalization between the PLA signal and OPP dense translation foci in UV-treated cells, suggesting that MATMs localize on γ-tubulin. Mean data on the right show increase in PLA signal (or colocalization between FTO and γ-tubulin) in UV treated cells compared to control cells. n=3 experiments, *p<0.0001 to control.

### MARK4 activates FTO in stress by phosphorylating threonine-6 (T6)

Using a demethylation assay, we measured FTO m6A demethylation activity, immunoprecipitated from UV or 2DG treated cells, and found an increase in its activity within 20 min of both UV and 2DG stresses **(Fig. 2A).** We predicted that FTO activity may be regulated by post-translational modifications, likely a phosphorylation tag, given the recognized role of kinase pathways in stress regulation^34, 35^. Using phospho-peptide enrichment and mass spectrometry of purified FTO from control and UV-treated cells, we found a phosphorylation tag at T6, induced by 20 min of UV stress **(Supp. Results 1, Table 1)**. Mutation of T6 to a phosphorylation mimetic (glutamic acid) or a non-phosphorylation mimetic (alanine) showed respectively increased and decreased FTO activity in vitro at baseline **(Fig. 2B)**, suggesting that the T6 phosphorylation is an important residue for regulation of FTO demethylation activity. We then generated and validated a polyclonal antibody specific for pT6-FTO. We found an increase in pT6-FTO levels with both UV and methionine deprivation treatments (**Figs. 2C-D**). pT6 levels from purified FTO (immunoprecipitated from UV treated cells) showed a clear pT6 signal which decreased with addition of phosphatase in the test tube (**Fig. S2A**). In addition, using a different stress, pT6-FTO levels decrease with FTO siRNA treatment in methionine deprived cells **(Fig. S2B)**. To identify the kinase mediating acute FTO phosphorylation and activation, we first immunoprecipitated cytosolic FTO, looked at its binding partners and found that several of the binding partners are related to the cytoskeleton (**Fig. S2C**), in keeping with the immunofluorescence data of Figure 1. Based on structural modeling using AlphaFold2 (AF2), we predicted that MARK4, a microtubule associated stress kinase that has been linked to important diseases along with FTO^36–41^, can bind FTO (**Fig. 2E-F, S2D-E)**. AF2 revealed that the sites of interaction are between MARK4 catalytic site and FTO’s hypermobile, or dynamic, arms (that include S173 and T6) supporting a “touch-and go” interaction, enough to phosphorylate pT6-FTO. To prove a direct and functional interaction, we added human recombinant FTO and human recombinant MARK4 in a test tube without the presence of cell lysates. We found increased pT6-FTO levels by immunoblots, confirmed with mass spectrometry, and increased FTO demethylation activity in a sequential demethylation activity assay, both of which were prevented by the MARK4 kinase inhibitor OTSSP167^42, 43^ (**Fig. 2G, Supp. Results 1, Table 2**). The T6-FTO phosphorylation in UV-treated cells was also inhibited by MARK4 siRNA (**Fig. 2H**). Lastly, knock down of MARK4 and FTO resulted in significant decrease in OPP translation foci seen in stress, suggesting that both are required to induce the HARP translation in MATMs (**Fig. 2I)**. Overall, these data suggest that the microtubule-associated MARK4 can directly phosphorylate FTO at T6 acutely in stress, increasing its activity, and activating translation within MATMs.

**Figure 2:**
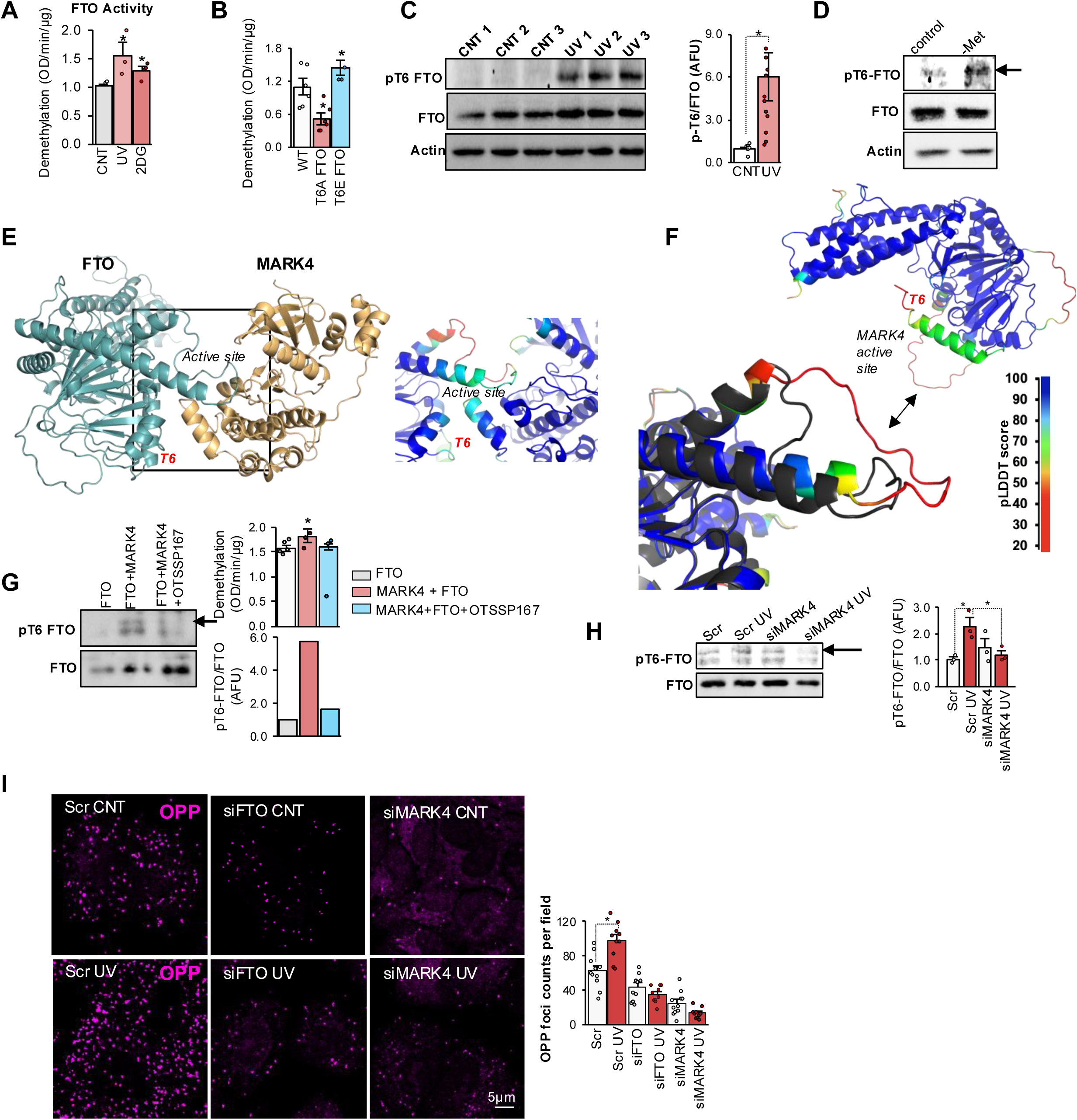
MARK4 activates FTO in stress by phosphorylating threonine-6. A) Mean data of percent increase in FTO activity measured using an m6A demethylase ELISA assay for immunoprecipitated FTO from UV or 2DG-treated cells. n=3 for UV, n=4 for 2DG, *p=0.018 and 0.006 compared to untreated control cells, respectively. B) FTO activity was measured at baseline using an m6A demethylase antibody-based assay of wildtype (WT), T6A, or T6E mutated FTO, overexpression in A549 cells and purified using flag-pull down. Demethylation activity decreases when T6 is mutated to alanine, a non-phosphorylated threonine mimetic, and increases when mutated to glutamic acid, a phosphorylated-threonine mimetic at baseline. n=6 for T6A, *p=0.014 compared to WT, and n=4 for T6E, p=0.017 to WT. C) Immunoblots showing increased FTO-T6 phosphorylation after 20 min of UV treatment detected using a custom phospho-T6 FTO specific polyclonal antibody from 3 different samples. Actin is used as a loading control. Mean data are shown on right. n=12, *p=0.03 compared to control (CNT). D) pT6-FTO and total FTO levels are measured using immunoblots in control and methionine deprived cells. Similar to UV, pT6-FTO levels increase with methionine deprivation stress. Full immunoblot including the effects of siFTO is shown in Fig. S2B. Actin is used as a loading control. E) Predicted structure of FTO (cyan) bound to MARK4 (aa 50-318, gold), generated using Alphafold2 (AF2). Residues 165-175 in FTO are predicted to bind MARK4 placing S173 directly into the active site of MARK4, marked “Active site”. Further contacts are predicted to occur in the helices formed by residues 36-46 in FTO, and 204-220 in MARK4. The T6-FTO region is within reach of the active site of MARK4 that includes a dynamic (mobile) arm. Black box indicates the interaction site. On the right is close-up image of the binding interface with residues of both proteins coloured by predicted Local Distance Difference Test (pLDDT) score. pLDDT scores greater than 90 (dark blue) indicate high confidence in both Cα and side chain positions, while scores between 70 and 90 indicate confidence in Cα structure alone (green-cyan) only. Scores below 50 indicate low confidence which is strong indicator of disorder or dynamic regions (red). The pLDDT score gradient is shown in Fig. 2F on the right. Segments of the loop directly involved in interacting with MARK4 are well defined with pLDDT scores ranging between 70 and 90, and the T6-FTO region is located is in proximity of the MARK4 active site. Note that the two loops (involving S173 and T6) that are within proximity of MARK4 active site are typically excluded in published FTO crystal structures due to their highly dynamic nature (e.g., PDB codes 8IT9, 3LFM, 4CXW, etc). F) Aligned loop region of bound FTO to MARK4 (grey) and unbound FTO residues (coloured by pLDDT scores). The disordered loop in the unbound form (with low pLDDT scores in red), is stabilized in the MARK4-bound FTO structure. *Above* is a predicted model of full length FTO with residues colored by pLDDT; the N-terminal helix is predicted to be flexible, and dynamic, specifically the first 6 residues including Threonine 6. The flexibility of the region suggests a kinase accessible region and a “touch-and-go interaction pattern. The other flexible region including S173 is in close proximity to the T6 flexible loop and the MARK4 active site. Double headed black arrow points to the same flexible loop in the full FTO model and the magnified overlaid model on the left. pLDDT score color legend is shown on right for both figures 2E and 2F. G) Recombinant MARK4 phosphorylates immunoprecipitated FTO from control cells at T6 in an in-vitro kinase assay (shown in immunoblots on left) and increases FTO demethylation activity, using an ELISA m6A demethylase assay (shown in chart on right). Both the T6 phosphorylation and the increase in the demethylation activity are decreased in the presence of the MARK 4 inhibitor OTSSSP167. n=5 for MARK4+FTO and n=3 for OTSSP167 group. *p=0.014 to FTO alone. The chart on bottom shows normalized levels of pT6-FTO/FTO of the displayed immunoblot. H) pT6-FTO levels after 20 min of UV decrease by pretreatment with MARK4 siRNA but not by scramble RNA. n=3, *p=0.008 for Scr and Scr UV and p=0.017 Scr UV and siMARK4 UV. I) Immunofluorescence staining of global translation measured using OPP (in magenta) in control and UV-treated cells in the presence and absence of FTO and MARK4 siRNAs. Mean data on right show a significant increase in translation foci after 20 min of UV stress in scrambled siRNA (Scr) only, but not in the presence of FTO or MARK4 siRNA. n=3, *p<0.05 compared to control (CNT).

### HARPs are synthesized within 20 min of stress from pre-existing mRNAs

To assess the overall degree of change in protein levels within 20 min of stress (UV and 2DG), we first utilized Stable Isotope Labelling with Amino acids in Culture (SILAC)^44^ followed by mass spectrometry, which allows for the grouping of proteins into increased, maintained, or decreased, by expressing protein levels as Heavy (stress, both UV and 2DG) over Light (control) ratios. While ∼70% of proteins remained unchanged, 25% decreased and ∼5% increased in stress (**Fig. 3A, Fig. S3A-B)**. RNA sequencing from the same cells revealed that the increased proteins did not correlate with their corresponding RNA levels, suggesting that the regulation is at the translation level and that some proteins can be synthesized within 20 min from stress from pre-existing mRNAs, in keeping with our hypothesis (**Fig. 3B)**. Although this gave us a bird’s eye view of the protein homeostasis in the first 20 min of stress, SILAC does not label exclusively the proteins synthesized *after* the induction of stress, due to the prolonged incubation of cells with the labelled amino acids over several passages. Thus, we also utilized L-AHA-labeled proteomics with mass spectrometry to characterize the newly synthesized proteins further and ensure that the increased levels are not due to decreased degradation. Cells were incubated with L-AHA immediately after exposure to stress for 20 min and using Click chemistry reaction on beads, newly synthesized proteins from control and UV treated cells were enriched and identified with mass spectrometry from three biological replicates. This method gave us similar results to SILAC with 5% of proteins increasing within 20 min from stress. To identify which of the increased proteins were synthesized in an FTO-dependent manner according to our hypothesis, we gave the FTO inhibitor DAC51^45^, or DAC, 30 min prior to stress and L-AHA labelling. We used a specific inhibitor as opposed to an FTO siRNA to prevent the confounding effects of long-term inhibition of FTO on RNA homeostasis, since siRNA takes much longer than 20 min to inhibit FTO. We defined HARPs as proteins that significantly increased within 20 min of UV compared to control cells, without an associated increase in their mRNA. ∼73% of HARPs were also responsive to some degree to DAC inhibition with a range between 20% and 100% inhibition, suggesting that FTO plays an important role in regulating translation of a large percentage of HARPs (**Fig. S3C)**. Treatment of cells with the eIF4G1 inhibitor SBI7556^46^, or SBI, which is a cap-dependent translation inhibitor, resulted in ∼50% decrease in both total proteins and HARPs, in contrast to DAC that decreased the levels of HARPs but not of total proteins, suggesting that DAC selectively inhibits HARPs translation without affecting the overall protein translation **(Fig. 3C)**. The AHA-labeled HARPs were decreased by both DAC and SBI, while the SILAC-labeled decreased proteins (i.e., the non-HARPs) were predictably decreased by UV without any significant DAC or SBI effects. These data suggest that FTO selectively promotes HARP translation and that many HARPs are translated in cap-dependent manner; but cannot rule out that some HARPs can also be translated by cap-independent translation pathways, a redundancy not surprising for stress response proteins that are critical for cell survival.

**Figure 3.**
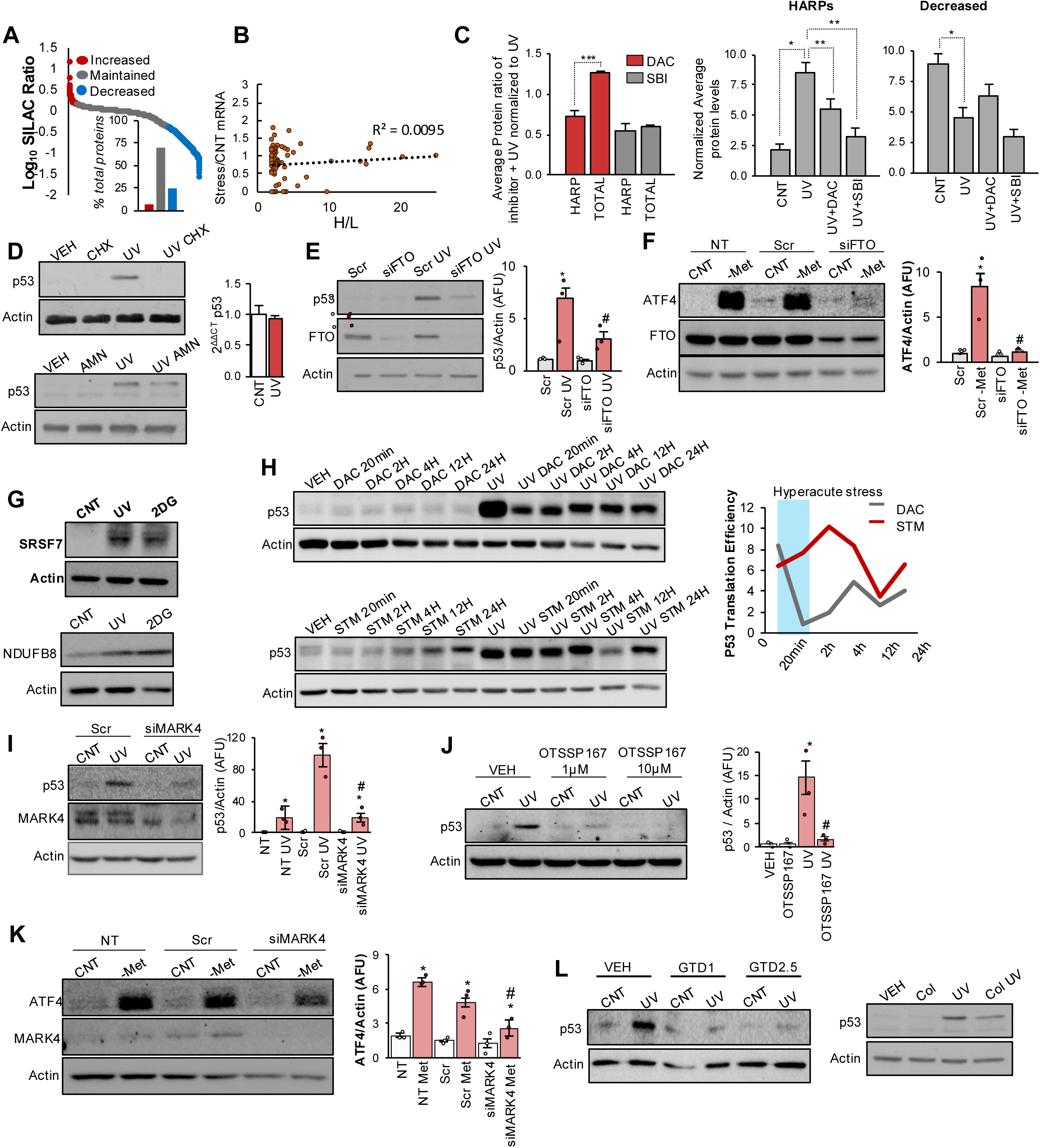
HARPs are synthesized within 20 min of stress from pre-existing mRNAs. A) Log_10_ of SILAC ratios (expressed as heavy (H) for UV and 2DG / light (L) for control) of all protein hits shown in decreasing order as a dot plot, from 4 biological replicates. Red and blue outline the increased and decreased hits, respectively, while the grey color represents the maintained, or relatively unchanged, hits. Increased hits are 213 distinct proteins, maintained are 2490, and decreased are 869. Cutoff points to separate the groups of proteins were selected based on natural breaks in data. Percentage of each protein group are shown in chart below. B) Correlation plot of RNA levels from SILAC experiment measured using RNA-seq, of the top increased hits in SILAC showing poor correlation between increased proteins and their RNA levels. Dotted line shows line of best fit of data points. C) HARP levels were detected in UV treated cells after AHA-incorporation using mass spec in presence and absence of DAC (DAC51, FTO inhibitor) treatment or SBI (SBI7556, an eIF4G1 inhibitor). Protein levels were normalized by ranking, and Welch’s t test is used to find the significantly increased proteins in the UV group compared to control group from n=3 experiments. Average fold change of protein levels (based on rank data of 3 biological replicates) between UV and UV + DAC or UV + SBI were calculated. HARPs are defined as proteins that are significantly increased with UV treatment, compared to control, within 20 min. The effect of FTO inhibitor, DAC, or cap-dependent translation inhibitor, SBI, on HARP translation or on total protein translation is in chart on left showing that FTO inhibition decreases only HARP translation but not total translation, whereas cap-dependent translation decreases both HARP and total translation equally. Average normalize protein levels for all 4 groups for HARPs and decreased proteins are outlined in middle and right charts, respectively. Full list of HARPs is shown in **Fig. S3C**. *p<0.05 control (CNT) to UV, **p<0.05 UV to UV+DAC or UV+SBI, ***p=0.00007 to TOTAL. D) p53 levels are measured 20 min after UV treatment in the presence and absence of CHX (cycloheximide, ribosome inhibitor), or AMN (a Amanitin, RNA Pol II inhibitor) using immunoblots at the top or bottom, respectively. Actin is used as loading control. VEH = vehicle. On right, chart shows p53 mRNA levels measured after UV, using qRT-PCR exhibiting no change in p53 mRNA levels within 20 min of UV. 18S was used as a loading control. CNT = control. E) p53 protein levels are measured after 20 min of UV treatment in the presence and absence of siRNA targeting FTO (siFTO) or Scrambled (Scr), using immunoblots. Actin is used as loading control. Mean levels of p53 protein normalized to Actin, are shown on the right. *p=0.003 compared to Scrambled control (Scr), #p=0.019 compared to Scr UV. n=3. F) ATF4 protein levels are measured after 30 min of methionine deprivation (-Met) in the presence and absence of siRNA for FTO (siFTO) using immunoblots. Actin is used as a loading control. Mean data of normalized ATF4 protein levels are on the right. n=3, *p=0.023 to control (non-Methionine deprived group), #p=0.024 to Scr -Met group. Scr = Scrambled siRNA. G) Immunoblots showing an increase in SRSF7 and NDUFB8 HARP proteins after 20 minutes of UV and 2DG. Actin is used as a loading control. H) p53 protein levels are measured after treatment time course with DAC51 (DAC), a potent and selective FTO inhibitor (top immunoblot), and STM2457 (STM) (bottom immunoblot), a potent and selective METTL3 inhibitor, for the times shown using immunoblots. Actin is used as a loading control. p53 translation efficiency (TE) was measured as a ratio of normalized p53 protein levels to actin over normalized p53 mRNA (2^DDCT^ to 18S). Effect on TE were seen as early as 20 minutes with DAC pre-treatment but not with STM. Blue bar highlights the 20min zone which is the timeframe of the studied stress model. I) Immunoblots showing p53 protein levels after UV treatment in the presence and absence of siRNA for MARK4. Actin is used as a loading control. CNT= Control, Scr = scrambled siRNA. n=3. Mean data are shown on right. *p=0.001 compared to control (Scr), #p=0.004 compared to Scr UV. J) Immunoblots showing p53 protein levels after UV treatment in the presence and absence of OTSSP167 at 1µM or 10µM. Actin is used as a loading control. VEH= vehicle. Mean data are shown on right. *p=0.019 compared to control (VEH), #p=0.024 compared to UV. n=3. K) ATF4 protein levels were measured after 30 min of methionine deprivation (-Met) in the presence and absence of siRNA for MARK4, using immunoblots. Actin was used as loading controls for immunoblots. n=3, *p=0.007 compared to non-deprived group. #p=0.027 compared to Scr -Met (scrambled methionine deprived) group. L) Immunoblots showing p53 protein levels in UV treated cells with presence or absence of gatastatin (an inhibitor of γ-tubulin GTPase activity) at 1µM or 2.5µM (GTD1 and GTD2.5) respectively, on the left, and colchicine (Col, a microtubule destabilizer) on the right. Actin is used as a loading control. VEH = Vehicle (DMSO).

The AHA-list of HARPs included several proteins that are well-known master regulators of the stress response like the transcription factors p53^47^ and ATF4^48^, but also several other proteins not previously linked to stress. Because transcription factors and particularly p53 have very low levels and are difficult to be reliably identified with mass spectrometry, we also used targeted mass spec to confirm its presence on in this list **(Supp. Results 1, Table 3**). To further validate the HARPs’ increase with immunoblots, we used methionine restriction for ATF4^48^ as inducing trigger (as ATF4 is known to be particularly responsive to this stress) and UV or 2DG for p53, SRSF7 and NDUFB8 from our HARP list. p53 levels increased with UV, and cycloheximide (a ribosome inhibitor) but not α amanitin (an RNA polymerase II inhibitor^49^), prevented p53 increase, while its mRNA levels did not increase (**Fig. 3D**). In addition, as p53 is known to also be regulated at the protein degradation level, time course with nutlin-3A, a specific MDM2 inhibitor^50^ which causes buildup of p53 levels, did not result in p53 protein levels increase to the level of UV (**Fig. S3D)**. siRNA for FTO prevented the acute increase of p53 with UV stress confirming that it is required for the acute translation of p53 (**Fig. 3E)**. In addition, overexpression of T6E (phospho-mimetic) FTO, but not T6A, resulted in increased p53 levels even in absence of UV stress **(Fig. S3E)**. These data show that p53 is translated acutely and is in fact a HARP and underly the importance of T6 phosphorylation on FTO activity and its role in HARP regulation under stress. ATF4 protein increased with methionine deprivation stress, but not at the mRNA level, and this was prevented by FTO siRNA (**Fig. 3F, Fig. S3F)**. The splicing factor SRSF7, which has recently been shown to be important for lung and colon cancer cell survival^51^ was also found to be increased within 20 min of both UV and 2DG stresses, as did NDUFB8 (**Fig 3G**). Altogether, these data provide evidence that HARPs are translated acutely within minutes of various stresses, either using AHA-labeling with mass spectrometry, or with immunoblots validation.

We then used DAC51 pre-treatment using increasing timepoints from 30 min to 24hr, to see at what point does FTO activity inhibition impact translation. This was also compared to STM2457^52^ (or STM), a METTL3 inhibitor, for the same timepoints. We found that FTO inhibition, as early as 30 min, inhibits the stress-induced increase in p53 translation efficiency (i.e., protein/mRNA levels), but not STM, which inhibits it around the 12-hour point **(Fig. 3H)**. This confirms that FTO acute activation, and the removal of an inhibitory m6A tag, is required for p53 translation.

In addition to A549 cells, the increased protein levels of p53 or SRSF7, within 20 min of UV stress in an FTO-dependent manner (using the FTO inhibitor DAC51) was reproduced in primary lung fibroblasts **(Fig. S3G)** and 3 additional cancer cell lines: A101D primary melanoma **(Fig. S4A)**, A2058 metastatic melanoma **(Fig. S4B)** and CCL227 metastatic colorectal cancer **(Fig. S4C)**. Furthermore, inhibition of MARK4 using siRNA or pre-treatment with the MARK4 small molecule inhibitor OTSSP167^43^ prevented the pT6-FTO and HARPs’ increase within 20 min of UV stress in A549 cells **(Fig. 3I-J),** or in A101D (**Fig S4A**), A2058 (**Fig S4B**), CCL227 (**Fig S4C**) and fibroblasts **(Fig. S3H).** Knockdown of MARK4 using siRNA also prevented ATF4 translation in methionine deprivation stress, **(Fig. 3K)**. ATF4 was also induced within 20 min of UV stress, and this was once again prevented by MARK4 siRNA **(Fig. S3I)**. Taken together these data show that HARPs can be induced in an FTO and MARK4-dependent manner in several cell types under diverse stresses, suggesting that the HARP/MARK4 axis may be a previously unrecognised stress response pathway.

Lastly, destabilization of microtubules with colchicine or inhibition of γ-tubulin GTPase activity with gatastatin^53^ 3 minutes prior to UV stress, also inhibited p53 translation (**Fig. L**), suggesting that MATMs are also required for HARP translation.

### Specialized cytoskeletal ribosomes are important for HARP synthesis in stress

Actively translating ribosomes, known as polysomes, are either free-floating in the cytosol or embedded in ER sheaths, with recent reports of ribosomal sub-pools localizing to mitochondrial membranes^14^. While our understanding of ribosomal heterogeneity is expanding^17, 19^, and few reports have suggested binding of ribosomes directly to microtubules, there has been no evidence to date of *active* ribosomes on microtubules that are distinct from ER ribosomes^16, 54, 55^. Our data suggested the presence of active ribosomes in stress within MATMs microdomains since MATMs hosted translation of peptides containing OPP or L-AHA. We used transmission electron microscopy (TEM) and phosphorus mapping^56^, which allows for the sensitive and specific detection of ribosomes, to visualize the microtubules and ribosomes with nanometer resolution. This demonstrated clear ribosomal particles at the microtubule junctions (i.e., γ-tubulin), highlighted within dotted circles in **Fig. 4A**. Regular TEM also showed three distinct pools of ribosomes (cytosolic, ER, and cytoskeletal-CS) (**Fig. 4B**).

**Figure 4:**
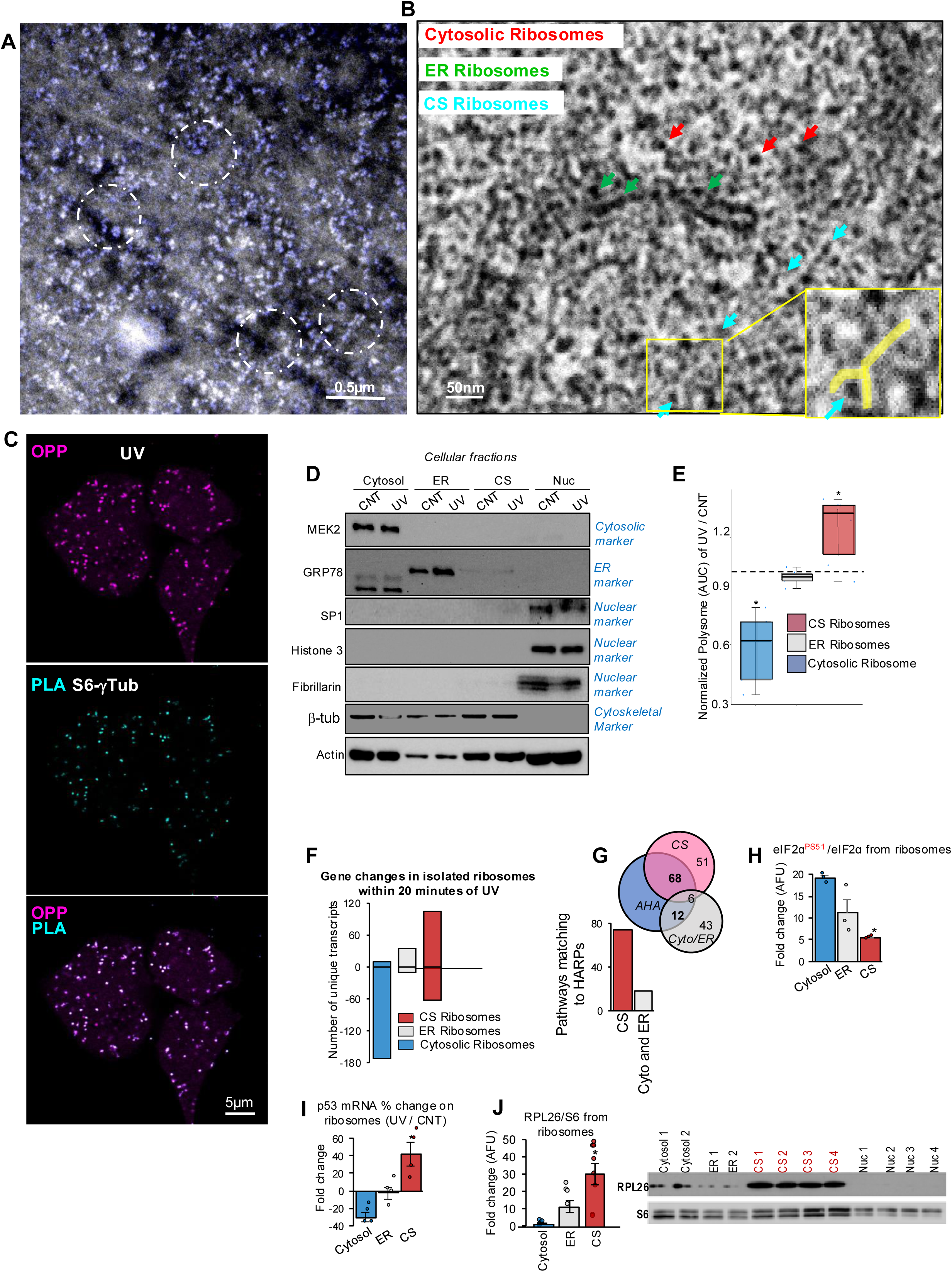
Specialized cytoskeletal Ribosomes are important for HARP synthesis in stress. A) Transmission electron microscopy with overlayed phosphorus mapping, which detects ribosomes (in blue) of A549 cells, showing ribosomes over microtubule branching points (characteristic of γ-tubulin), with examples highlighted within white dotted circles. B) Transmission electron microscopy (TEM) imaging of UV-treated A549 cells showing ribosomes in the cytosol (red arrow), ER sheets (green arrow), and along microtubule (blue arrow). CS ribosomes can be seen at microtubule junction “Y” forks as shown in blue arrows. Inset shows a magnification of region enclosed in yellow square. “Y” fork (reminiscent of g tubulin) is highlighted in yellow within the magnified inset. C) Immunofluorescence staining of OPP (in magenta) and PLA between S6 and γ-tubulin (in teal) in UV-treated cells showing colocalization between the PLA signal and OPP translation foci. D) Immunoblots showing fractionation purity of cytosolic, ER, cytoskeletal (CS), and nuclear fractions, in control (CNT) and UV treated cells. Subcellular fraction markers are shown in blue font on the right; Cytosolic marker: MEK2, ER marker: GRP78, Cytoskeletal (microtubules) marker: β-tubulin, Nuclear markers: SP1, H3, Fibrillarin. E) Area under the curve of polysomal peaks representing active translation obtained using HPLC of ribosomes (Mega-SEC) of UV-treated cells normalized to control from different cellular fractions: Cytosolic, ER, and Cytoskeletal (CS). Controls and absorbance peaks are shown in Fig. S6A-B. Data show increased polysomal AUC indicating increased translation within 20 min of UV stress in CS ribosomes, with decreased AUC and translation in Cytosolic and relatively unchanged AUC in ER ribosomes. n=5, *p=0.0006 and *p=0.004 for Cytosol and CS Ribosomes compared to ER Ribosomes, respectively. F) Ribosome sequencing of ribosome-protected mRNA fragments in Cytosol, ER, and CS isolated ribosomes, measured by differential sequencing analysis between UV and control treated cells within each ribosomal group, showing only the statistically significant differentially expressed hits within each pool. The largest number of increased translated hits within 20 min of UV stress occurs in the CS ribosomes, and the largest number of decreased hits is in the Cytosolic ribosomal fraction after 20 minutes of UV. n=3 biological replicates. G) Number of pathways of the significantly increased transcripts in the Cytoskeletal (CS) ribosomes and pooled cytosolic and ER ribosomes (Cyto/ER) of Ribosome sequencing, that detects and quantifies ribosome-bound mRNAs, matching to AHA pathways of increased genes detected in L-AHA mass spectrometry quantification of the newly translated proteins (Fig. 1I). Venn diagram at the top shows total number of pathways from CS and Cyto/ER ribosomes, and the bottom chart shows only the number of pathways matched to HARP pathways from AHA experiment. H) Mean data of pS51-eIF2a levels normalized to total eIF2a measured using immunoblots (sample blot shown in Fig. S5D) in Cytosol, ER, or Cytoskeletal (CS) isolated ribosomes in UV-treated cells. Data show significantly lower pS51-eIF2a levels in the CS ribosomes (suggesting increased translation) compared to either Cytosol, or ER ribosomes post UV stress. n=3, *p<0.05 compared to cytosol and ER ribosomes. I) p53 mRNA levels measured as fold change in UV-treated cells compared to control cells in Cytosol, ER, or CS isolated ribosomes using qRT-PCR. 18S is used as a loading control. n=4, *p=0.001 and p=0.013 compared to Cytosol and ER groups, respectively. J) Immunoblot example of A549 cells and mean data of RPL26 protein levels normalized to S6 ribosomal protein levels measured using immunoblots in Cytosol, ER, or CS isolated ribosomes in UV treated cells. Data show RPL26 enrichment in the CS ribosomes compared to ER or cytosolic ribosomes. n=6, *p=0.0001 and p=0.008 compared to Cytosol and ER ribosomes, respectively.

Since we see a strong correlation between FTO, γ-tubulin and OPP signal acutely in stress **(Fig. 1H)**, we predicted that PLA between S6 ribosomal protein and the γ-tubulin/OPP signal, would result in same overlap signal, which is exactly what we found **(Fig. 4C)**. In addition to A549 cells, we found a similar overlap in 3 additional cell lines (A2058, A101D, CCL227), and primary lung fibroblasts), shown in **(Fig. S5A-B**) suggesting that CS ribosomes are present in several cancer and normal cell lines.

To better assess active translation functionally, we isolated the ribosomal pools from cytosolic (Cyto), ER, and cytoskeletal (CS) fractions by modifying an established protocol to fractionate cells and isolate ribosomes from different fractions^15^. We verified the separation of fractions using markers for cytosol (MEK2), ER (Grp78), β-Tubulin for cytoskeleton, and SP1, H3, Fibrillarin for nuclear fractions **(Fig. 4D)**. We utilized ribosomal profiling using HPLC^57^ to quantify polysomes at baseline and UV stress by measuring the area under curve (AUC) of absorbance peak at 260nm from polysomes (**Fig. S6A).** Immunoblots from different fractions corresponding to time on the HPLC column, show the ribosomal proteins corresponding to the absorbance peaks (**Fig. S6B)**. Polysomal AUC ratio of stress to control revealed decreased translation in the cytosolic fraction, unchanged translation in the ER fraction, and increased translation in the CS fraction within 20 minutes of UV **(Fig. 4E)**. We then performed ribosome-sequencing (Ribo-seq) where only the mRNA fragments protected by ribosomes are sequenced ^58^. Using Differential Gene Expression analysis of mRNA hits, where only significantly different hits are represented, we found that cytosolic ribosomes had the greatest number of decreased bound mRNA after 20 minutes of UV, while the ER ribosomal pool showed little change and the CS ribosomal pool had the largest differential increase in number of transcripts bound to ribosomes after stress **(Fig. 4F)**. Intriguingly, we also saw a significant number of proteins that decreased in the CS fraction as well, suggesting that CS ribosomes may have alternate translation roles in addition to HARP translation in stress. Pathway analysis of the proteins that are either differentially upregulated in each ribosomal pool or were specific to each ribosomal pool (labeled as CS enriched, or Cytosol and ER enriched) based on Ribo-seq, revealed that the CS-enriched proteins matched closer the pathways detected in the AHA labelling mass spectrometry experiments, suggesting that the newly translated HARPs within 20 minutes of UV are more closely linked to the CS ribosomal pool **(Fig. 4G)**.

p-eIF2α levels, (a translation initiation factor that decreases translation rates when phosphorylated at S51 in stress ^59–61^), were highest in the cytosolic ribosomal fraction, and lowest in the CS ribosomal fraction 20 minutes post UV **(Fig. 4H, Fig. S5C)**. To characterize the CS ribosomes further, we utilized mass spectrometry analysis of isolated ribosomes proteome from the three extra-nuclear cell fractions and found that the CS-ribosomal proteins and interactome (which includes associated proteins, nascent proteins, ribosomal proteins) clustered together using hierarchal clustering (**Fig. S6C**) with significant differences from the ER ribosomes. Pathway analysis of the significant proteins are shown in **Fig. S6D.** Important differences included translation regulation proteins such as IF4A1, which is a helicase associated with aggressive ER negative breast cancer with ability to bind RNA Quadruplexes (RG4) ^62^, IF4A3, which is involved in RNA splicing ^63, 64^, along with other translation and stress regulators, which are discussed in more detail in the ***Supplementary Results 2***. Lastly, we found that p53 mRNA increases significantly in CS ribosomes, compared to other ribosomal pools 20 min post UV (**Fig. 4I)**. Further supporting their structural differences from ER ribosomes, CS ribosomes were enriched in RPL26, that has been shown to be important for p53 translation ^47^ **(Fig. 4J)**. These data suggest the presence of functionally and structurally distinct ribosomes as integral parts of MATMs where compartmentalized HARP translation takes place in hyper-acute stress.

### A unique m6A signature marks HARP mRNAs

The dependence on FTO activation suggests that m6A demethylation allows HARP translation by demethylating and removing inhibitory m6A tags off their mRNAs, which are known to supress translation^7–9^. According to our hypothesis, the pre-stress (baseline) m6A signature may allow recognition of HARP mRNAs as they exit the nucleus, to be transferred on MATMs for translation at a later point. HARP recognition may involve a unique methylation pattern and because m6A is so abundant, a single base resolution of m6A locations is required to detect a pattern of methylation unique to HAPRs. Thus, we investigated published miCLIP m6A databases which provide single base resolution of m6A peaks in A549 cells at baseline^65^. We normalized the m6A location by mRNA length to be able to compare the methylation patterns and we then grouped the m6A locations by protein levels in stress from our AHA experiment into increased (i.e., HAPRs), maintained and decreased protein groups. We found that compared to non-HARP mRNAs, HARP mRNAs had a specific pattern of m6A methylation characterized by concentration of the peaks along the middle of the mRNA as two large peaks (**Fig. 5A)**. In addition, the mean peak location, in reference to 5’UTR, was significantly higher (or more downstream) in the HARP and decreased protein groups compared to the maintained group (**Fig. 5A-B)**. The HARP m6A pattern seemed to have more in common with the decreased compared to the maintained protein groups. However, the decreased proteins had more m6A peaks along the 3’UTR, unlike HARPs which had the majority of their peaks along the exons (**Fig. 5B-C)**. Note that in Fig. 5B, the mean location of m6A peaks in the HARP group is significantly more downstream (from 5’UTR) as compared to the Maintained protein group, suggesting that HARP m6A peaks are more likely to be in middle of the transcript, overlapping exons, unlike Maintained and Decreased transcripts, which tend to be more upstream (closer to 5’UTR), and overlapping 3’UTR, respectively. Although this pattern is perhaps not specific enough, it at least suggests that HARP mRNAs may be recognized for transport separately from non-HARP mRNAs although it is possible that additional recognition sites may exist in the 2- or 3-dimentional conformation of HARP mRNAs.

**Figure 5:**
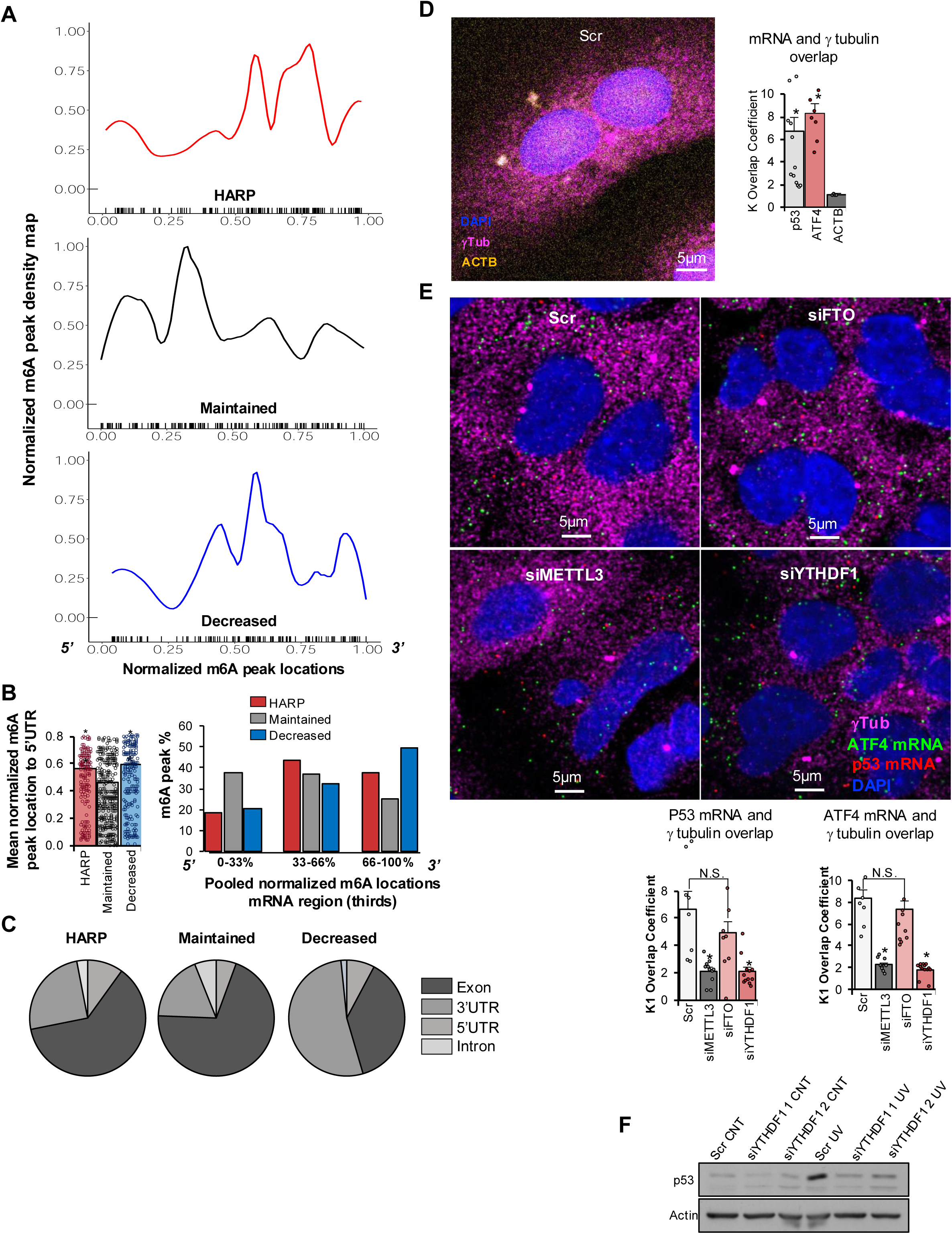
A unique m6A signature marks HARP mRNAs. A) Density plots of pooled A549 miCLIP m6A peaks normalized to mRNA length, grouped based on protein translation change with UV into HARP (Increased proteins within 20 min of UV stress), Maintained, or Decreased, as per mass spectrometry labelling of AHA tagged peptides (Fig. 3C). X-axis 0 represents 5’ and 1 represents 3’ of mRNA. The curve line represents the mean density of peaks, the grey regions around the curve lines represent the peak location variability. Individual peaks for each group are shown as black lines along the x-axis. B) On the left, mean normalized m6A peaks location per protein group are shown. Maintained proteins have lower mean location of m6A peaks, or more upstream, as compared to HARP (or increased) and decreased proteins, which have a higher, or more downstream, mean m6A peak location. *p<0.00001 compared to Maintained. Shown on the right are mean normalized m6A peak location per protein group divided into 3 equal segments from 5’ to 3’ of the mRNA transcript, showing that HARP mRNAs have more peaks in the middle region of the transcript (where exons are more likely to be located) compared to Maintained and Decreased proteins. The individual peaks and their locations are shown in Fig. 5A. C) Pie charts of labelled mRNA features where m6A peaks are found per protein group, HARP, Maintained, and Decreased. Decreased proteins have the majority of the m6A peaks in 3’UTR, unlike Maintained, or HARP (or increased) proteins. Note that in B, the mean location of m6A peak of HARP group was significantly more downstream as compared to Maintained, suggesting that HARP m6A peaks are more likely to be in middle of the transcript, overlapping exons, unlike Maintained and Decreased transcripts, which tend to be more upstream, or overlapping 3’UTR, respectively. D) Immunofluorescence staining of β-Actin mRNA (in yellow), γ-tubulin (magenta) and DAPI (blue). Mean data of K1 overlap coefficients between p53 (red), ATF4 (green), β-Actin (ACTB) and γ-tubulin in control cells are shown on right. n=4 *p=0.024 and p=0.002 for p53 and ATF4 compared to ACTB, respectively. E) Immunofluorescence staining of p53 mRNA (in red), ATF4 mRNA (in green), γ-tubulin (in magenta), DAPI (in blue), in Scrambled (Scr), or siRNA against FTO, METTL3, or YTHDF-treated cells. Mean data of K1 overlap coefficients between p53 (red), ATF4 (green), and γ-tubulin in cells treated with siRNA targeting FTO, METTL3, or YTHDF1 are shown below. p<0.05 compared to Scrambled siRNA (Scr). n= 9-14 fields. *p=0.003 to CNT for siMETTL3 and p=0.0001 for siYTHDF1 for p53 mRNA. *p<0.0001 for siMETTL3 and siYTHDF1 for ATF4 mRNA. F) p53 levels were measured after UV treatment in the presence and absence of 2 siRNAs targeting YTHDF1, or Scrambled (Scr) siRNA, using immunoblots. Actin was used as loading control.

We then looked at m6A readers and focused on the well-studied YTHDF1^66, 67^ speculating that it may be involved in the recognition of HARPs and directly or indirectly in their transportation to MATMs at baseline. We used siRNA to METTL3 (to inhibit the m6A writer), to FTO, (to inhibit the m6A eraser) and to YTHDF1 (to inhibit a reader) on non-stressed A549 cells, in addition to mRNA hybridization for HARP mRNAs (i.e. p53, ATF4) and non-HARPs (e.g. actin) and studied their colocalization with MATMs (γ-tubulin). We found that compared to non-HARP, HARP mRNAs were localized more with γ-tubulin, compared to actin (**Fig. 5D)**. The colocalization within MATMs (marked by OPP foci and γ-tubulin) in stress, decreased with inhibition of YTHDF1 and METTL3 but not with FTO inhibition (**Fig**. **5E****, Fig. S7A**). These data suggest that METTL3 is required to add the methylation tags for HARP mRNAs, and YTHDF1 is required to read them and is likely involved in directing HARP mRNAs to MATMs. In contrast, FTO’s decrease by siRNA at baseline (when it has not yet been activated by MARK4) does not have an impact on HARP mRNA localization; FTO is only needed after stress to demethylate m6A and allow its translation. Knockdown of YTHDF1 with siRNA also prevented p53 acute translation (**Fig. 5F)**. Taken together these findings suggest that m6A likely “marks” HARP mRNA with a specific pattern of methylation that may be involved in their suppression of translation at baseline and in their transport to MATMs, where activated FTO may allow their translation to proceed upon stress, while the translation of non-HARPs outside of MATMs remains inhibited during stress.

### Disrupting the HARP synthesis system decreases survival after UV and chemotherapies

If the hyperacute synthesis of HARPs is critical for stress defence and survival, inhibition of FTO or its activator at stress (i.e., MARK4) should lead to increased death. FTO inhibition with siRNA led to a significant increase in apoptosis, measured by cleaved caspase-3, 24 hours after a UV pulse (**Fig. 6A**). Pre-treatment of A549 cells for 20 min prior to UV with the FTO inhibitor DAC, caused a significant increase in UV-induced death, measured by propidium iodide 24 hrs post UV, compared to cells pretreated with vehicle (**Fig. 6B**). We assessed apoptosis at 24 hours (and not 20 minutes) because the earliest time point to detect caspases with immunoblots is at least 24 hours. However, the cell’s commitment to enter apoptosis versus adaptation pathways (like the HARP pathway) happens quickly after stress, perhaps within 20 minutes. To better understand the timing of the FTO-driven survival benefit, we placed cells under a HoloMonitor microscope, which can measure growth and cell confluence by continuous imaging based on optical density signals of unstained cells, thus avoiding any additional stress from live straining. The DAC-treated cells had significantly decreased confluence at 36 hours compared to cells exposed to UV alone, and this was obvious quite early post stress (**Fig. 6C**). The MARK4 inhibitor OTSSP limited the UV-induced pT6-FTO levels (**Fig. 6D**), limited the large p53 protein induction 20 min post UV (**Fig. 6D**), mimicking as expected the effects of FTO inhibition (see Fig. 3E) and significantly increased apoptosis at 24 hours (**Fig. 6E**).

**Figure 6:**
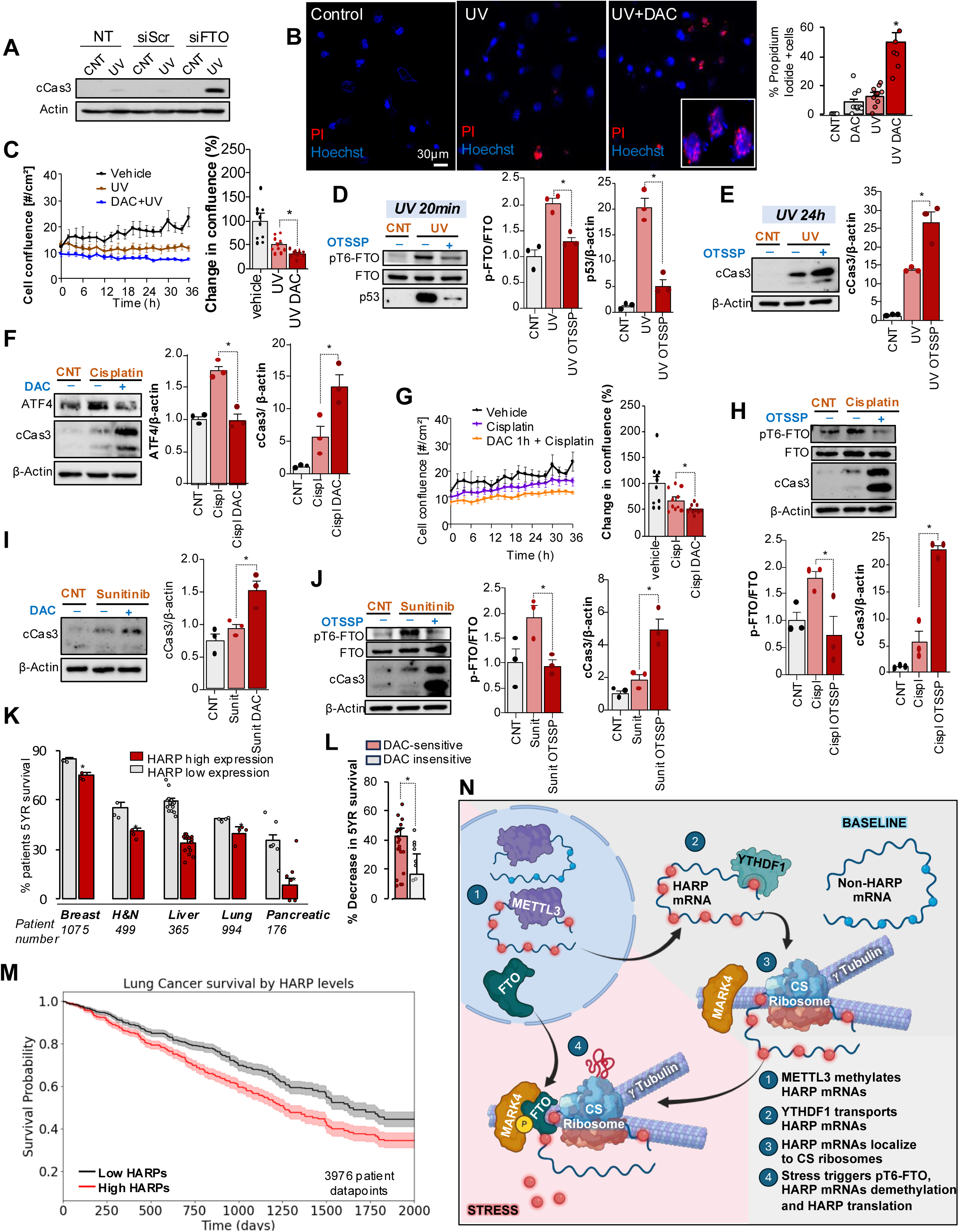
Disrupting the HARP synthesis pathway decreases survival after stress. A) Apoptosis measured by cleaved caspase 3 (cCas3) levels 24 hours after UV treatment in non-transfected and Scrambled (Scr) or FTO siRNA transfected A549 cells. Actin is used as a loading control. B) Propidium Iodide (PI, a marker of cell death) staining of A549 cells 24h after treatment with UV or UV + DAC. Hoechst, blue, marks the nuclei. Inset in bottom right shows higher magnification of PI overlapping with nuclei. Mean data and SEM are shown right. n=5-7, *p=0.00001 to UV. C) Cell confluence measured using live continuous imaging with A HoloMonitor microscope in cells treated with UV or UV + DAC over 36 hours. Mean data with SEM are shown on right. n=3, *p=0.0002 compared to UV. D) pT6-FTO levels, along with total FTO and p53 in UV and UV+OTSSP (MARK4 inhibitor, applied 2 min prior to UV)-treated cells. Mean data with SEM are shown on right for pT6-FTO normalized to FTO and p53 levels normalized to Actin. n=3, *p=0.0021 and p=0.001 compared to UV for normalized p-FTO and p53 respectively. E) Apoptosis measured by cleaved caspase-3 levels in UV or UV+OTSSP-treated cells for total treatment duration of 24h using immunoblots. Mean data normalized to Actin are shown on right. n=3, *p=0.007 compared to UV. F) ATF4 and cCas3 levels measured using immunoblots in Cisplatin or Cisplatin+DAC treated cells using immunoblots. Actin is used as a loading control. Mean data with SEM for normalized ATF4 and cCas3 levels are shown on right. n=3, *p=0.002 and p=0.017 compared to Cisplatin (Cispl) for normalized ATF4 and cCas3, respectively. G) Cell confluence measured with a HoloMonitor microscope in Cisplatin or Cisplatin+DAC-treated cells over 36 hours. Mean data with SEM are shown on right. n=3, *p=0.011 compared to Cisplatin. H) pT6-FTO levels, along with total FTO and cCas3 levels measured in Cisplatin, or Cisplatin+OTSSP-treated cells using immunoblots. Actin is used as loading control. Mean data with SEM for normalized pT6-FTO and cCas3 levels are shown below. n=3, *p=0.025 and p=0.0007 compared to Cisplatin for normalized p-FTO and cCas3, respective. I) cCas3 levels are measured in cells treated with Sunitinib or Sunitinib+DAC using immunoblots. Actin is used as loading control. Mean data with SEM are shown on the right. n=3, *p=0.013 compared to Sunitinib. J) pT6-FTO levels, along with total FTO and cCas3 levels are measured in Sunitinib or Sunitinib+OTSSP-treated cells using immunoblots. Actin is used as loading control. Mean data with SEM for normalized pT6-FTO and cCas3 levels are shown on the right. n=3, *p=0.01 and p=0.007 compared to Sunitinib for normalized p=FTO and cCas3, respectively. K) % 5-year survival of patients based on tumor samples from various cancers (breast, head and neck, liver, lung, pancreas) in 176-1075 patient samples from the RNA-sequencing and immunohistochemistry database in *Protein Atlas*, based on high versus low expression of HARPs (shown in Fig S3C). There was significant survival benefit for patients with low HARP-expressing tumors (p<0.001), suggesting higher resistance to stress in tumors expressing high levels of HARPs. Treatment status of each sample is unknown but because tumor excision and biopsy occur prior to chemotherapy, these samples are typically at baseline prior to chemotherapy. The average decrease in 5-year survival for all HARPs was measured at 35% (*p<0.05 to low HARP expression for each cancer type), which is of significant magnitude in Oncology standards. L) Mean % decrease in 5-year survival from the Protein Atlas database among all of the 5 tumors types studied 5 cancer types sub-grouped into DAC-sensitive (>20% decrease in HARP translation with DAC) and DAC insensitive (<20% decrease in HARP translation with DAC, see Fig 3C, S3C) showing that DAC-sensitive HARPs had a greater prognostic value for overall survival; *p<0.05. M) Kaplan Meier curve of survival probability of lung cancer patients divided by level of HARP expression. High HARP (shown in red) patient group shows decreased survival compared to low expression group (shown in black). N) Proposed pathway for a comprehensive cap-dependent translation of HARM mRNAs in Microtubule Associated Translation microdomains (MATMs) on γ-tubulin. In addition to the m6A decoration that non-HARP mRNAs have (depicted by blue dots), HARP mRNAs have an additional m6A signature (depicted by red dots), which allows them to be recognized by m6A readers like YTHDF1 and transferred to MATMs, which host specialized ribosomes, whereas the non-HARP mRNAs are transferred to canonical ribosomes. Upon stress, FTO exits the nucleus and is localized to MATMs. There, the microtubule associated kinase MARK4 phosphorylates FTO on T6, activating its demethylase activity. This removes the m6A HARP signature, accelerating the HARP mRNA translation rate, whereas the non-HARP translation rate remains suppressed outside of MATMs.

Our hypothesis suggested that the synthesis of HARPs may be important in all stresses and our mechanism was studied in the UV, 2DG and methionine restriction stresses. We speculated that HARPs and FTO activation will also be important in perhaps the biggest stress that a cancer cell can be exposed to, i.e. chemotherapy. We thus tested the effects of FTO and MARK4 inhibition on cells exposed to either a genotoxic chemotherapy (cisplatin) or a more targeted agent (sunitinib). DAC prevented the cisplatin-induced increase in the HARP ATF4 (**Fig. 6F**). This was associated with a decrease in confluency at 36 hours (**Fig. 6G**) similar to what we found in UV. Most importantly OTSSP decreased pT6-FTO levels and increased apoptosis 24 hrs post either cisplatin or sunitinib (**Figs. 6H-J**). These data suggest that FTO activation contributes to cancer cells defense against stress, regardless of its type. It also suggests that FTO and particularly MARK4 inhibitors can be used along with chemotherapy (or radiation therapy) to increase the effectiveness of therapy.

Our list of HARPs was discovered in an unbiased manner and included both known and novel stress response proteins. We studied our list of HARPs (shown in **Fig. S3C**) against the public database of the Protein Atlas Project^68^ (www.proteinatlas.org), where the level of specific tumor proteins measured with RNA sequencing and immunohistochemistry (“high vs low” protein levels), are correlated to 5-year survival data from hundreds to thousands of patients with different cancers. We found that patients with tumors that had high levels of our HARPs had on average 35% decrease in 5-year survival, compared to patients with tumors having lower HARP levels. This was seen in breast, head and neck, liver, lung, and pancreatic cancers, with each patient cohort having survival data from hundreds of patients **(Fig. 6K)**. Importantly, the decrease in survival was mostly driven by HARPs whose synthesis was inhibited by DAC with levels of inhibition ranging between 20 and 100% (i.e., DAC-sensitive), compared to an inhibition of 0-20% (i.e., DAC-insensitive) (**Fig. 6L)**. Survival plots for individual cancers, with pooled HARPs data for tumor-high versus tumor-low low expression of HARPs showed worse survival for patients with high HARP levels in liver cancer (**Fig. 6M)**, and lung, breast, pancreatic cancers, i.e., the tumors with the highest number of patients and HARPs available in the Protein Atlas database **(Fig. S7C)**. These data support our hypothesis that HARPs enhance the survival of cancer cells to stress and offer an indirect but intriguing independent validation of the importance of the HARP synthesis pathway in cancer patient survival, potentially in multiple cancers.

## Discussion

Here we describe a comprehensive pathway by which cancer cells can synthesize HARPs immediately after diverse stresses (including the stress of chemotherapy) from pre-existing mRNAs, that appears to coordinate their stress response, since inhibition of this pathway suppresses HARP synthesis and increases their vulnerability to stress, and subsequent death. Our proposed mechanism is summarized in (**Fig. 6N)**: Upon exit from the nucleus, m6A-decorated HARP mRNAs are transferred to translation microdomains (MATMs) on γ-tubulin, that host specialized ribosomes. While their m6A decoration keeps their translation suppressed at baseline, MARK4 activates FTO during stress (which exits the nucleus immediately after stress) which in turn demethylates HARP mRNAs, allowing their selective translation during stress, while non-HARP mRNA translation, outside of MATMs, remains suppressed. This is facilitated by a specific pattern of m6A decoration on HARP mRNAs (shown by the red tag, over and above their background m6A decoration pattern), which is distinct from the non-HARP mRNAs (shown by the blue tags, **Fig. 6N**). The idea that stress response can be coordinated by a set of proteins synthesized within 20 min from stress is new and may lead to a new approach in preventing the ability of cancer cells to resist and adapt to stress, a major challenge in oncology.

Following an unbiased proteomics and transcriptomics approach we discovered proteins that have not so far been implicated in stress response, perhaps due to our focus on the hyperacute stage of stress, that is not often followed in stress biology. This list (Fig S3C) also included well-known stress response proteins like the master transcription factors p53 and ATF4. For the extensively studied p53, we showed that the increase in its protein levels within 20 min from stress is not due to inhibition of degradation from the proteasome, a common assumption, but is a part of the acute synthesis of HARPs. The same is true for ATF4, the translation of which is presumed to be due to uORF and eIF2α phosphorylation. Both p53 and ATF4 were inhibited by cytosolic FTO inhibition. It is likely that for such central coordinators of the stress response like p53 and ATF4, redundant pathways exist for their translation, both cap-dependent and cap-independent. This is supported by recent evidence that has linked m6A to ATF4 translation which has canonically been known to be regulated by uORF alternative translation and eIF2α phosphorylation^69^. Both p53 and ATF4 have many non-transcriptional functions and interactions with stress pathway proteins, which may be more critical for their hyperacute effects, compared to their longer-term transcriptional effects.

Our work suggests that inhibiting the HARP synthesis comprehensively may target a previously unidentified Achille’s heel in cancer’s early stress adaptation processes. Indeed, while our discovery of HARPs was unbiased, it turned out that many of our HARP protein levels are associated with worse 5-year survival in many cancers, supported by the published survival data in the cancer *Protein Atlas*. Our data are also in keeping with the promising effects of FTO inhibitors in recent cancer trials, and particularly their ability to suppress cancer stem cells, the ultimate stress resistant cells^45, 70, 71^. However, MARK4 inhibitors many be more specific, since they can disrupt the HARP synthesis by inhibiting cytoplasmic FTO at stress (since MARK4 is a microtubule associated kinase), sparing nuclear FTO and its many known (and unknown) demethylase effects in the nucleus.

Another important aspect of this work is that it can explain how HARP synthesis is activated during stress, where translation inhibition is well established^60, 61, 72^. This is achieved by a compartmentalization of HARP translation in MATMs, in keeping with the increasingly recognized role of compartmentalization of translation in cell biology^73^. To the best of our knowledge this is the first time that functional specialized ribosomes are described on the cytoskeleton (γ-tubulin).

Our data are also in keeping with the increasingly recognized role of the m6A axis in cancer progression and resistance^11^. While METTL3^74^ and YTHF1^75–77^ have been directly implicated, less is known about FTO, and particularly cytosolic FTO. Our finding that FTO exits the nucleus immediately after stress, suggest that FTO’s ability to inhibit HARP synthesis is based on its location in the cytoplasm and MATMS, at least during the hyperacute stage of stress. A potential role of demethylating other unknown but potentially relevant targets beyond m6A cannot be ruled out, considering the many cytoskeletal proteins that we found interacting with FTO (Fig S2C).

Beyond FTO, this work suggests that the other components of our proposed pathway can be used in the fight against cancer’s adaptive response to stress, like γ-tubulin and MARK4. Our work is the first to implicate γ-tubulin in mRNA translation, the inhibition of which by gatastatin also prevented the stress-induced increase in HARP synthesis (Fig. 3L). While more work is needed to elucidate the role of γ-tubulin and its GTPase on HARP mRNA translation and the stress response in caner, our work is in keeping with the fact that increased γ-tubulin levels are associated with increased cancer aggressiveness and decreased survival in many cancers^28, 78–80^. Similarly, this is the first time that the role of MARK4 is shown in the direct activation of FTO in the cytoplasm. MARK4 is not studied well in cancer although some papers have shown the ability of MARK4 inhibitors in increasing the susceptibility of hepatocellular carcinoma to paclitaxel^81^. We propose that FTO, MARK4 and γ-tubulin inhibitors can all be used in association with cancer therapies against cancer resistance. We also propose that T6-FTO can be an important biomarker for cancer resistance before or during cancer therapy.

**Limitations:** More work is needed in answering the following three questions:

1. What is the mechanism for FTO’s exit from the nucleus? FTO is known to shuttle between the nucleus and cytoplasm and it is possible that this may be due to a phosphorylation as FTO can be phosphorylated at multiple locations with diverse effects in both its location and demethylation activity. Indeed, a recent paper showed that phosphorylation of nuclear FTO at T150 by nuclear casein kinase II, controls its nuclear retention/exit^82^, but remains unknown whether nuclear casein is responsive to stress. Along with our finding that MARK4 phosphorylation at T6-FTO, increasing its demethylation activity, these findings suggest that double FTO mutants on T6 and T150, maybe a useful tool with which to study the spatiotemporal regulation of FTO in stress in the future.
2. What is the precise nature of the activation signal for MARK4? Many kinases can be sensitized by other stress kinases. An example of an upstream kinase is GSK3, an important metabolic regulator ^83^ that is likely to play a role at least in our metabolic stress models. Thus, it remains to be determined whether MARK4 activation is the primary or secondary stress sensor in one or some of our stress models and whether the mechanism for its activation is phosphorylation by another stress kinase or another stress-sensitive post translational modification taking place within 20 min of stress.
3. What is the precise HARP m6A signature and its role? The pattern of m6A methylation that we identified in HARPs (Fig 5A-C), may not be specific enough to be called a “signature” although it was based on published miCLIP data that have single base resolution. More importantly, is the role of such an m6A signature to a) offer the well-known translation inhibition (so that HARPs are not synthesized and induce unnecessary reprogramming in the absence of stress, b) offer a recognition pattern for m6A readers that can facilitate HARP mRNA transfer to MATMs, or both? Are there one or two signatures that can allow those two functions in the 2 dimensional or 3-dimensional conformation of HARP mRNAs? We believe that our identification of specific HARPs can facilitate the discovery of a putative HARP mRNA signature in the future.

Overall, this work may open a new window in the fight against cancer adaptation to stress with major implications for oncology (Fig 6) and may also shed more light in stress biology in general, since we found that at least some aspects of the HARP pathway are also operational in non-cancer cells like fibroblasts (Fig. S3F-G, S5A-B).

## Supplementary material

### Supplementary Results

#### Supplementary Result 1: Mass Spectrometry tables

**Table 1.** p-T6 FTO peptide detected using phospho-peptide enrichment and mass spectrometry from UV treated cells.

| Peptide | threonine<br>modification | observed<br>m/z | MW<br>(exp.) | MW<br>(cacl.) | DeltaM<br>(ppm) |
| --- | --- | --- | --- | --- | --- |
| TPT<sup>6</sup>AEE<br>R | T6-Phosphorylation | 442.1816 | 882.3486 | 882.3484 | 0.28 |

**Table 2.** p-T6 FTO peptide detected using phospho-peptide enrichment and mass from immunoprecipitated FTO combined with recombinant MARK4.

| Protein<br>Accession | Descriptio<br>n | #PSMs | sequence | modification<br>s | charge | MH+ | DeltaM<br>(ppm) |
| --- | --- | --- | --- | --- | --- | --- | --- |
| Q9C0B1 | FTO | 140 | TPTAEE<br>R | T6-phospho | 2+ | 883.3574 | 1.82 |

**Table 3.** p53 peptide from UV treated cells, confirmed by targeted Mas-spectrometry.

| Annotated Sequence | m/z [Da] | MH+ [Da] | Group |
| --- | --- | --- | --- |
| [R].LGFLHSGTAK.[S] | 515.7895 | 1030.572 | control |
| [K].TcPVQLWVDSTPPPGTR.[V] | 955.9751 | 1910.943 | UV |
| [K].TcPVQLWVDSTPPPGTR.[V] | 637.6525 | 1910.943 | DAC+UV |

#### Supplementary Result 2: Pathway analysis of significant genes in CS ribosomes compared to ER ribosomes

##### CS ribosomes enriched GO pathways

*- RNA related:* Metabolism of RNA and regulation of RNA splicing, RNA localization.
*- Ribosome-related pathways:* ribonucleoprotein biogenesis and organization.
*- Cytoskeleton related pathways:* Non-membrane-bounded organelle assembly, regulation of actin cytoskeleton.
*- Other pathways:* nucleocytoplasmic transport, and regulation of cellular localization, among others.

These pathways suggest that composition of CS ribosomes and their interactome show specialized roles in RNA processing and regulation, Ribosomal biogenesis and regulation, cellular localization and transport, among others. Moreover, analysis of the decreased hits between ER and CS ribosomes show decrease in mitochondrial ribosomal proteins. This suggests that mitochondria are not enriched in the CS ribosomal pool, and that the CS ribosomes are unlikely to be associated with the previously described mitochondrial-associated ER ribosomes.

A closer look at the significantly increased proteins shows important roles in various pathways relating to stress translation. These include:

- *Translation control:* IF4A1, which is required for cap-dependent translation, IF4A3, which is an RNA helicase involved especially in RNA splicing. NOC4L, which is involved in selective mRNA translation, DHX9, which is involved in unwinding mRNA and initiation of translation, among others.
- *Stress pathways:* e.g. MTREX, which unwinds RNA duplexes and plays a role in DNA damage. SQTM and NKRF, or NFKb repressor factor, are involved in NFKB signaling and stress-related ribosomal processing respectively. TADBP, which associates with stress granules, and SUMO4, which binds to stress-defense proteins after oxidative stress.
- *RNA:* ROA2, which binds to m6A containing miRNA, FBRL, a methyltransferase involved in rRNA processing, HNRPC and B4DY08 which bind to poly-U tracts in 3’ or 5’ UTRs, along with various other mRNA splicing factors, such as SRSF7 and RNA helicases.
- *Ribosome binding proteins:* GAR1, involved in ribosome biogenesis and telomere maintenance, LASIL2 which is involved in biogenesis, among others.
- *Cytoskeletal proteins*: ITB1, involved in lamin matrix and cell motility, CKAP4 which mediates anchoring of ER to microtubules, and VIME which is involved in cytoskeletal stabilization and cell migration. ACTS and ACTG, which form Actin filaments, and PLEC which interlinks intermediate filamins and microtubules.
- *Localization:* RAB5C, which allows for docking and fusion of vesicles, CLH1, a vesicular protein, and F5H6E2, which is an unconventional myosin protein for intracellular movements, and F8VPF3, which is an ATPase cellular motor protein

##### CS ribosomes decreased GO pathways

While the majority of pathways that are significantly different between ER and CS ribosomal fractions were increased in CS ribosomes, a few pathways were significantly decreased including eIF3 complex pathway, regulation of translation pathway, mitochondrial ribosome, and mitochondrial translation initiation pathways.

---

**Figure S1.**
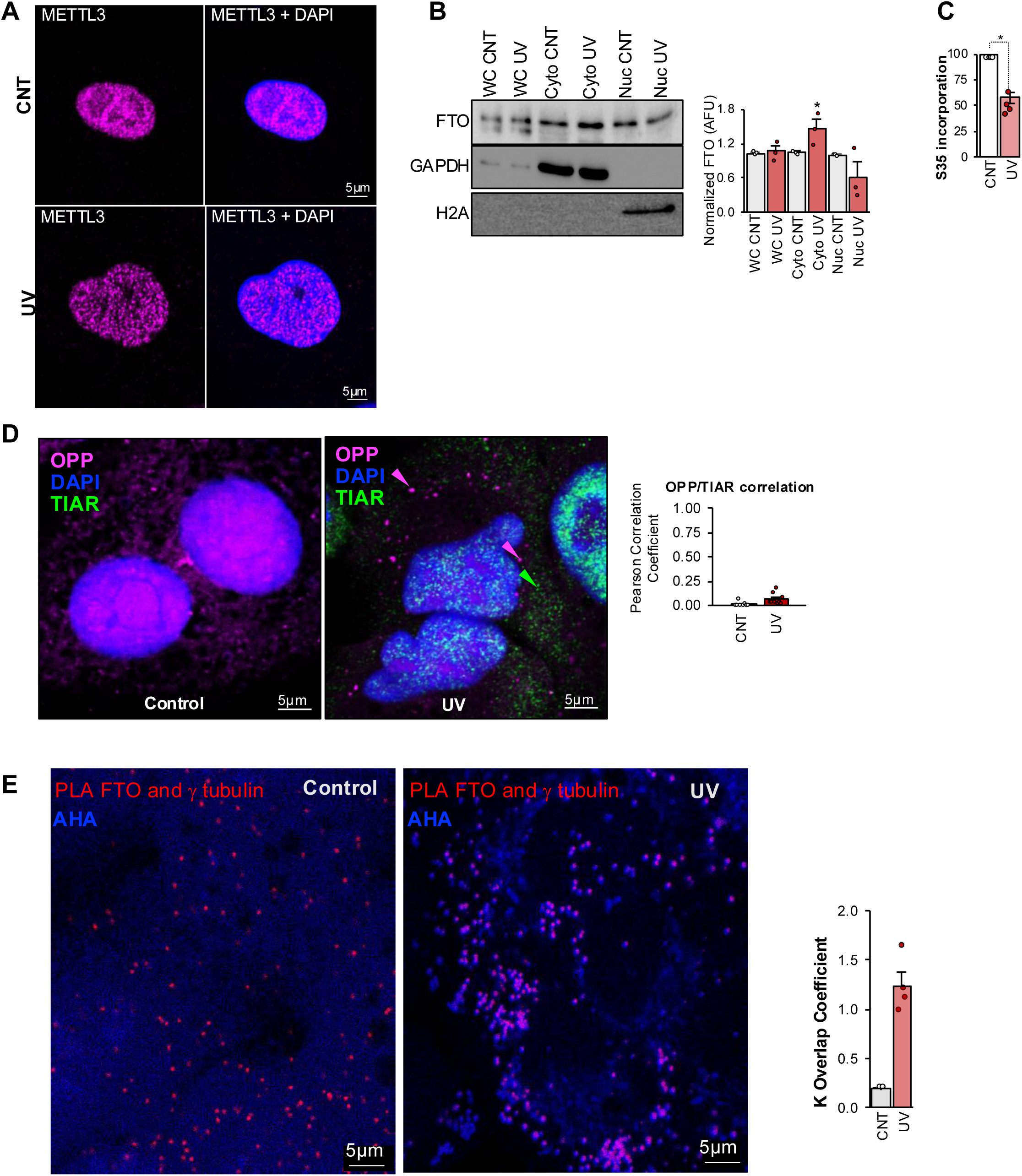
A) Immunofluorescence staining of METTL3 (magenta) and DAPI (blue) in control (CNT) and UV-treated cells showing an unchanged and exclusively nuclear signal for METTL3 in control and UV-treated cells. This representative image is from the experiments shown in Fig 1A, staining for METTL3 instead of FTO. B) Immunoblots showing FTO levels in whole cell (WC), Cytosolic (Cyto), and Nuclear (Nuc) fractions in control and UV treated A549 cells. GAPDH is used as a loading control for WC and Cyto, and histone 2A (H2A) is used as a loading control for Nuc fraction. Mean data for normalized FTO levels (to GAPDH for WC and Cyto, and to H2A (for Nuc) show increased FTO levels in Cytosol within 20 min of UV. p<0.05 compared to cyto CNT, n=3. C) To measure global translation after stress, newly translated proteins were radioactively labelled with Methionine S35 for 20 min in control and UV (25 J/m^2^ dose) treated cells and were run on SDS PAGE. Mean data of S35 signal, detected on radioactive-sensitive film, normalized to total protein Ponceau signal and to control (CNT) levels. Mean +/- SEM data are shown, n=3, *p<0.05 compared to control. The level of decrease in total protein translation was similar between radioactive S35 labelling and OPP staining, shown in Fig. 1B, within 20 min of UV stress. D) Immunofluorescence staining of OPP (magenta), DAPI (blue), and the TIAR stress granule marker (green) in control and UV-treated cells showing no colocalization between the dense translation foci (highlighted with purple arrows) induced after UV and the stress granule marker TIAR (highlighted with green arrow) that are also increased acutely in stress. Mean data of Pearson’s correlation coefficient between TIAR signal and OPP signal is shown for CNT and showing no correlation. n=8-9 fields. E) Immunofluorescence of control and UV-treated cells showing L-azidohomoalanine (AHA) labelling of newly synthesized proteins in blue along with Proximity Ligation Assay (PLA) staining of FTO and γ-tubulin in red. Mean data of overlap coefficient between PLA and OPP in control and UV treated cells is shown on right. n=4 experiments, *p=0.0002 to control.

**Figure S2.**
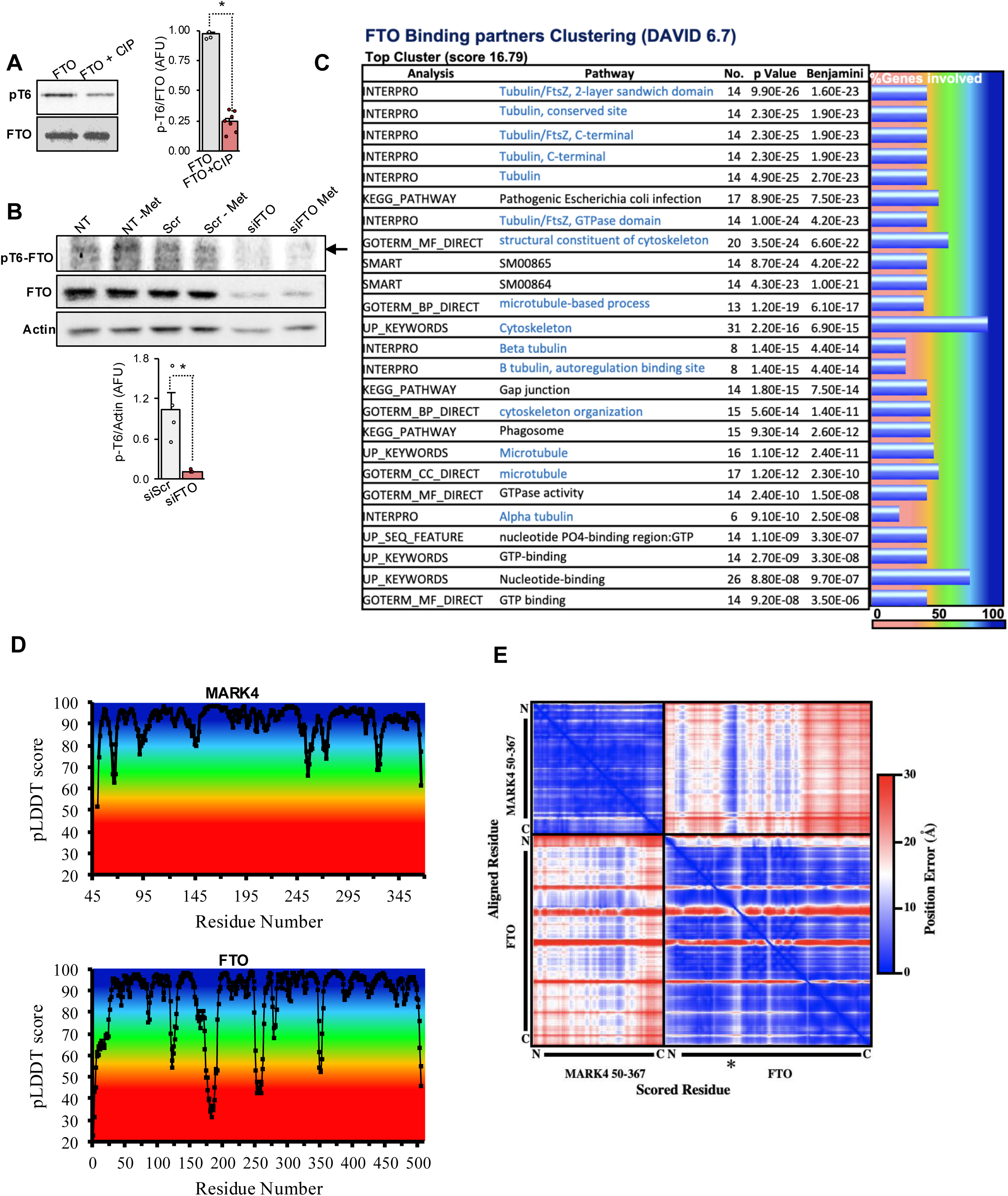
A) pT6-FTO and total FTO levels are measured using immunoblots in purified FTO (immunoprecipitated from UV treated cells) in phosphatase assay with Calf Intestinal alkaline phosphatase (CIP). Phosphatase decreases pT6-FTO levels, indicating specificity of pT6-FTO antibody. B) Immunoblot showing pT6-FTO levels, total FTO levels, with Actin as a loading control in cells treated with methionine deprivation in cells transfected with Scramble siRNA (siScr), or FTO siRNA (siFTO). Black arrow points to pT6-FTO band. siFTO shows decrease in both FTO and pT6-FTO levels, normalized to Actin. Mean data are shown below. p<0.05 compared to siScr. C) Chart showing FTO binding partners detected using FTO immunoprecipitation (at baseline) and mass spectrometry followed by clustering analysis using DAVID 6.7. Microtubule-related proteins are highlighted in blue-colored font. Nucleotide binding pathway was second most involved pathway. Blue bars on right side show the relative percentage of gene hits represented in the clusters indicated along a gradient scale shown at the bottom. D) pLDDT plot of MARK4 (Top) and pLDDT plot of FTO (Bottom). pLDDT scores are scaled from 0-100 and scores greater than 70 indicate high confidence in Cɑ position and scores greater than 90 indicate high confidence inside chain atoms. Typically scores lower than 50 are indicators of low confidence and suggest disorder. The background colouring is consistent with the colour scheme in figures 2 E-F. Overall the models have high pLDDT scores. FTO’s n-terminal residues were predicted with less confidence represented by pLDDT scores lower than 50, compared to other regions, suggesting a disordered or dynamic region. This pattern was also seen in the loop regions connecting helices and beta strands. E) Plot of predicted alignment error (pAE), also denoted as frame aligned point error (FAPE), for the predicted structure of MARK4 and FTO. Unlike pLDDT which is a local confidence metric, pAE is a long-range domain position confidence measurement calculated for each residue against all other residues. pAE is calculated as the predicted error in Angstroms (Å), at residue x if the model were aligned to the “true” structure at residue y. pAE does not require the residues to be from the same monomer, therefore it is practical to measure and analyze both intra- and inter-position confidence. The independent proteins were predicted with relatively high confidence, with FTO in the bottom right panel having a median error of 5.1 Å, and MARK4 in the top-left panel with a median error of 3.6 Å. Interestingly the error between MARK4 and FTO is relatively high in certain regions, but in regions denoted with asterisk (**\***), where the FTO loop is docked into the MARK4 active site, the median error is 6.1 Å, seen in the top right panel. This is the lowest predicted error between FTO and MARK4 suggesting a relatively confident model for interaction between FTO and MARK4 at those residues.

**Figure S3.**
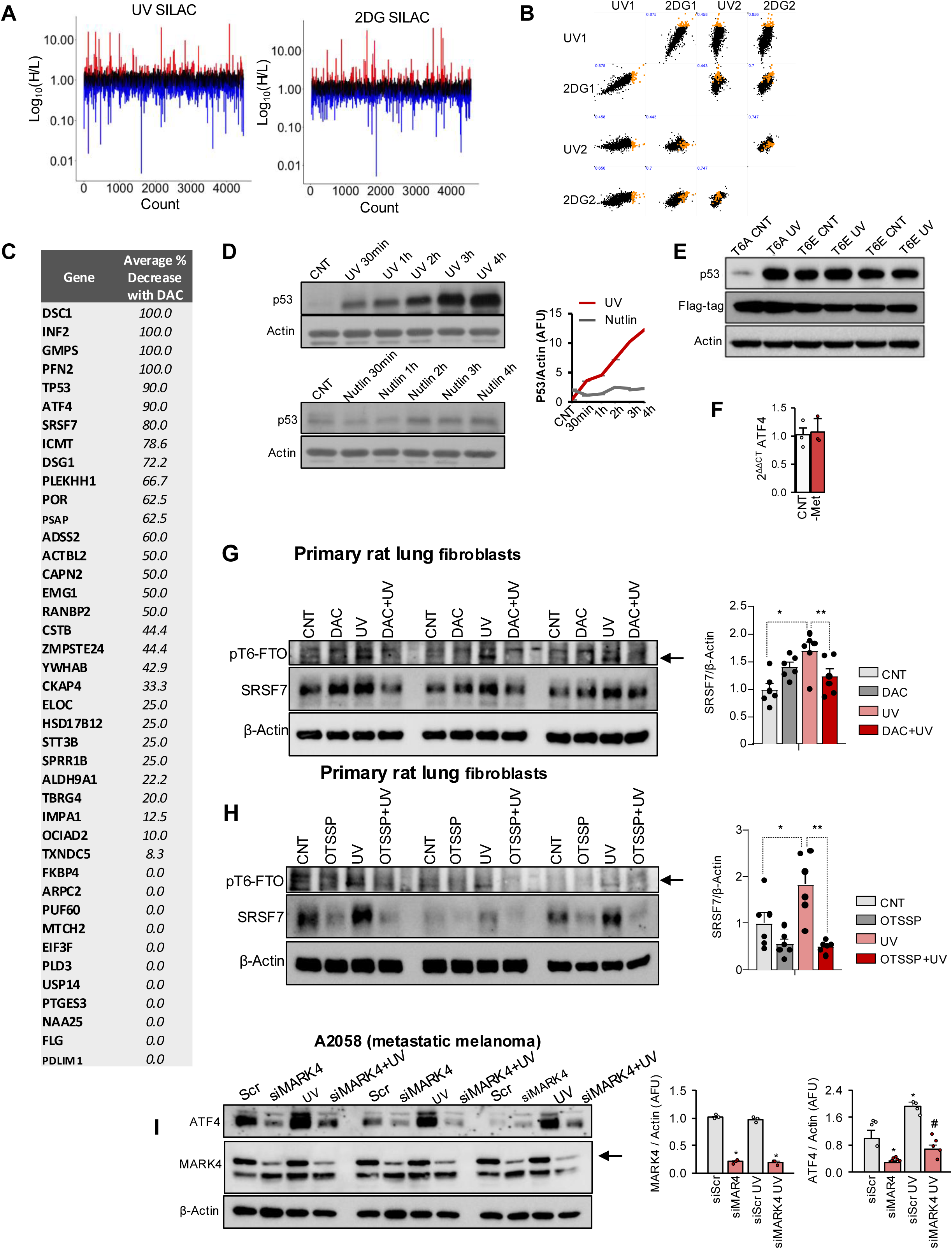
A) Line plot of total protein levels in UV (on left) and 2DG (on right), both Heavy labelled, compared to control Light labelled) cells, expressed as H/L ratio. Red lines are increased proteins, black lines are unchanged proteins, and blue lines are decreased proteins. B) Scatter plot analysis of protein levels in four stress SILAC groups, UV and 2DG, normalized to their controls as H/L ratios. Orange color is highlighting the “increased” protein groups in all 4 groups. C) HARPs list detected using AHA labelling, i.e., proteins that increased within 20 min of UV treatment, including HARPs like p53 or ATF4 or SRSF7 (also detected with SILAC) all of which were validated using immunoblots (in Fig. 3). D) Immunoblots showing p53 protein levels with a time course of UV (top) or Nutlin3A (Nutlin) treatment from 0 min to 4h with Actin as a loading control. Line chart on right shows the normalized p53 levels to Actin. E) Immunoblots showing p53 levels in T6A or T6E overexpressing A549 cells treated with UV. FTO is flag tagged, detected with anti-flag antibody. Actin is used as a loading control. F) ATF4 mRNA levels measured after Methionine deprivation, using qRT-PCR showing no change in ATF4 mRNA levels within 30 min of methionine deprivation. 18S was used as a loading control. CNT = control. G) Immunoblots showing pT6-FTO and SRSF7 protein levels with β-Actin as a loading control from primary lung fibroblasts treated with UV and DAC51. Cells were pre-treated with DAC 2 min prior to UV treatment. Mean data from 6 experiments are on the right showing increased SRSF7, normalized to Actin, with UV treatment which is inhibited with DAC treatment. Three biological replicates are shown on one gel. *p<0.05 control compared to UV. **p<0.05 UV compared to UV+DAC. H) Immunoblots showing pT6-FTO and SRSF7 protein levels with β-Actin as a loading control from primary lung fibroblasts treated with UV and OTSSP (or OTSSP167, MARK4 inhibitor). Cells were pre-treated with OTSSP, 2 min prior to UV treatment. Mean data from 6 experiments are on the right showing increased SRSF7, normalized to Actin, with UV treatment, which is inhibited with OTSSP treatment. Three biological replicates are shown on one gel. *p<0.05 control compared to UV. **p<0.05 UV compared to UV+OTSSP I) Immunoblots showing ATF4 and MARK4 protein levels with β-Actin as a loading control from A2058 metastatic melanoma cells treated with UV in presence and absence of siRNA for MARK4 (siMARK4). Cells were pre-treated with either scrambled siRNA or MARK4 siRNA for 2 days prior to UV treatment. Mean data from 5 experiments on the right show increased ATF4 protein levels normalized to Actin, with UV treatment, which is inhibited with siMARK4 treatment. Chart on left shows mean data for normalized MARK4 (n=3) with significant decrease in its levels with siRNA. Three biological replicates are shown on one gel. *p<0.05 siScr compared to siScr-UV. #p<0.05 siScr-UV compared to UV+siMARK4.

**Figure S4.**
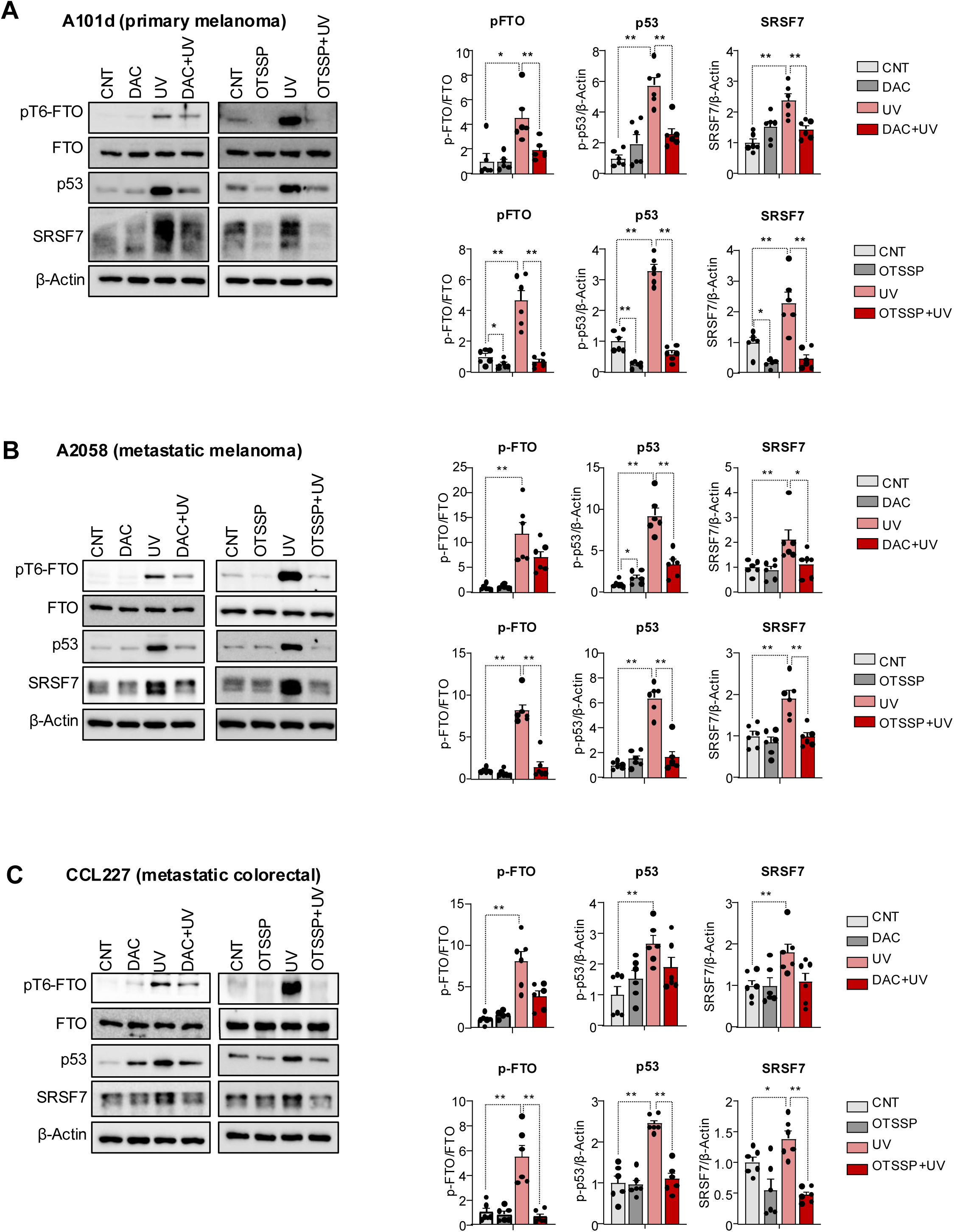
A) Immunoblots showing pT6-FTO, FTO, p53, SRSF7, and β-Actin as a loading control from A101d (primary melanoma cells) treated with UV and OTSSP (or OTSSP167, MARK4 inhibitor) on the left, or UV and DAC (DAC51, an FTO inhibitor). Cells were pre-treated with OTSSP or DAC, 2 min prior to UV treatment. Mean data from 6 experiments on the right show increased p-T6-FTO, p53, and SRSF7 normalized to Actin, with UV treatment, which are decreased with either OTSSP or DAC pre-treatment. B) Immunoblots showing pT6-FTO, FTO, p53, SRSF7, and β-Actin as a loading control from A2058 (metastatic melanoma cells) treated with UV and OTSSP (or OTSSP167, MARK4 inhibitor) on the left, or UV and DAC. Cells were pre-treated with OTSSP or DAC, 2 min prior to UV treatment. Mean data from 6 experiments on the right show increased p-T6-FTO, p53, and SRSF7 normalized to Actin, with UV treatment, which are decreased with either OTSSP or DAC pre-treatment. C) Immunoblots showing pT6-FTO, FTO, p53, SRSF7, and β-Actin as a loading control from CCL227 (metastatic colorectal, or CRC, cells) treated with UV and OTSSP on the left, or UV and DAC. Cells were pre-treated with OTSSP or DAC, 2 min prior to UV treatment. Mean data from 6 experiments on the right show increased pT6-FTO, p53, and SRSF7 normalized to Actin, with UV treatment, which are decreased with either OTSSP or DAC pre-treatment.

**Fig S5.**
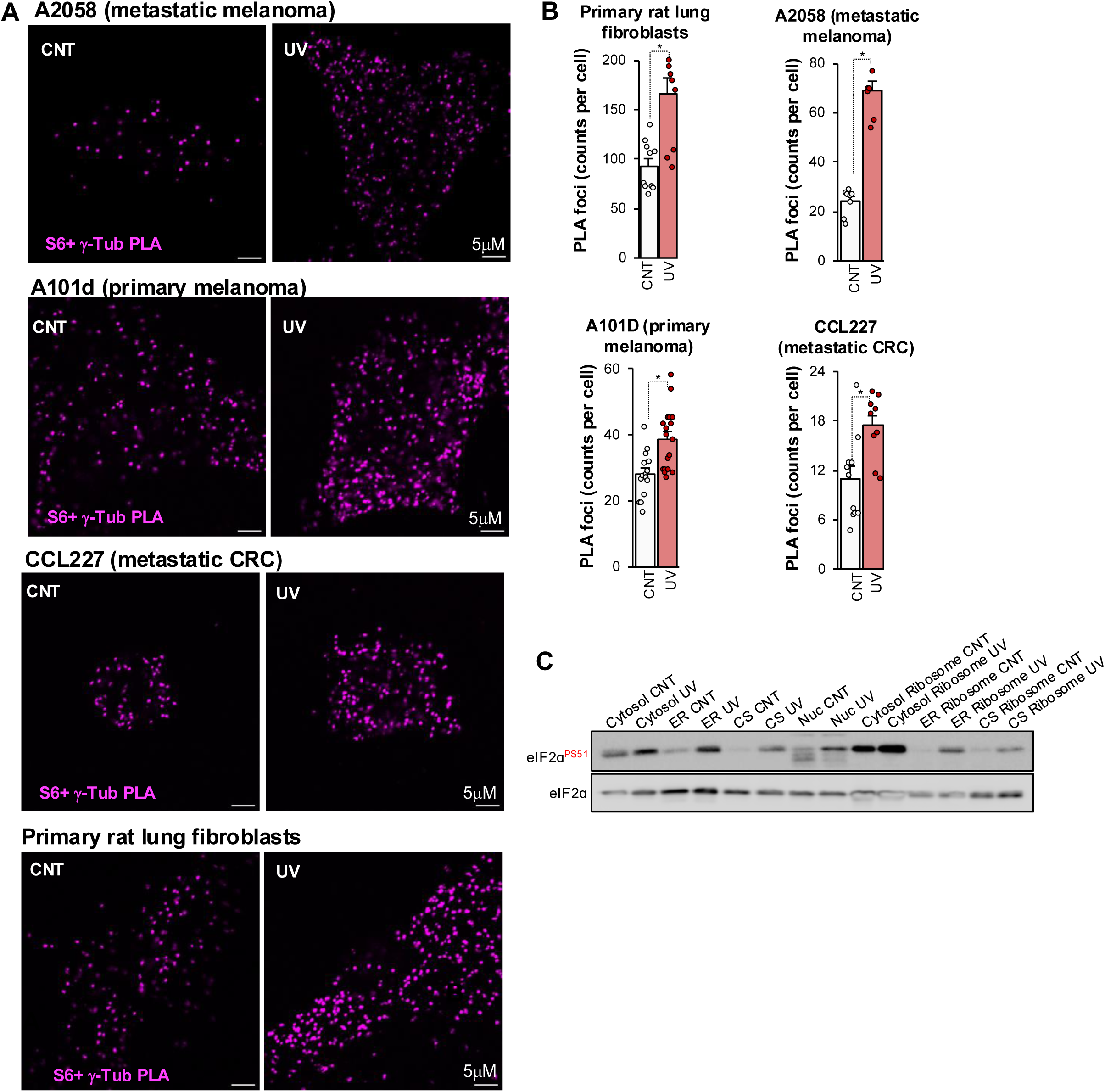
A) Representative images of S6 and γ-tubulin PLA signal in 4 lines in in control and UV-treated cells showing an acute increase in colocalization of S6 and γ-tubulin, after 20 min of UV exposure. B) Mean data from the experiments shown in A. *p<0.05 compared to CNT, n=9-17 fields from 3 biological replicates. C) Immunoblot from cellular subfractions and corresponding isolated ribosomes from those fractions showing total and

**Figure S6.**
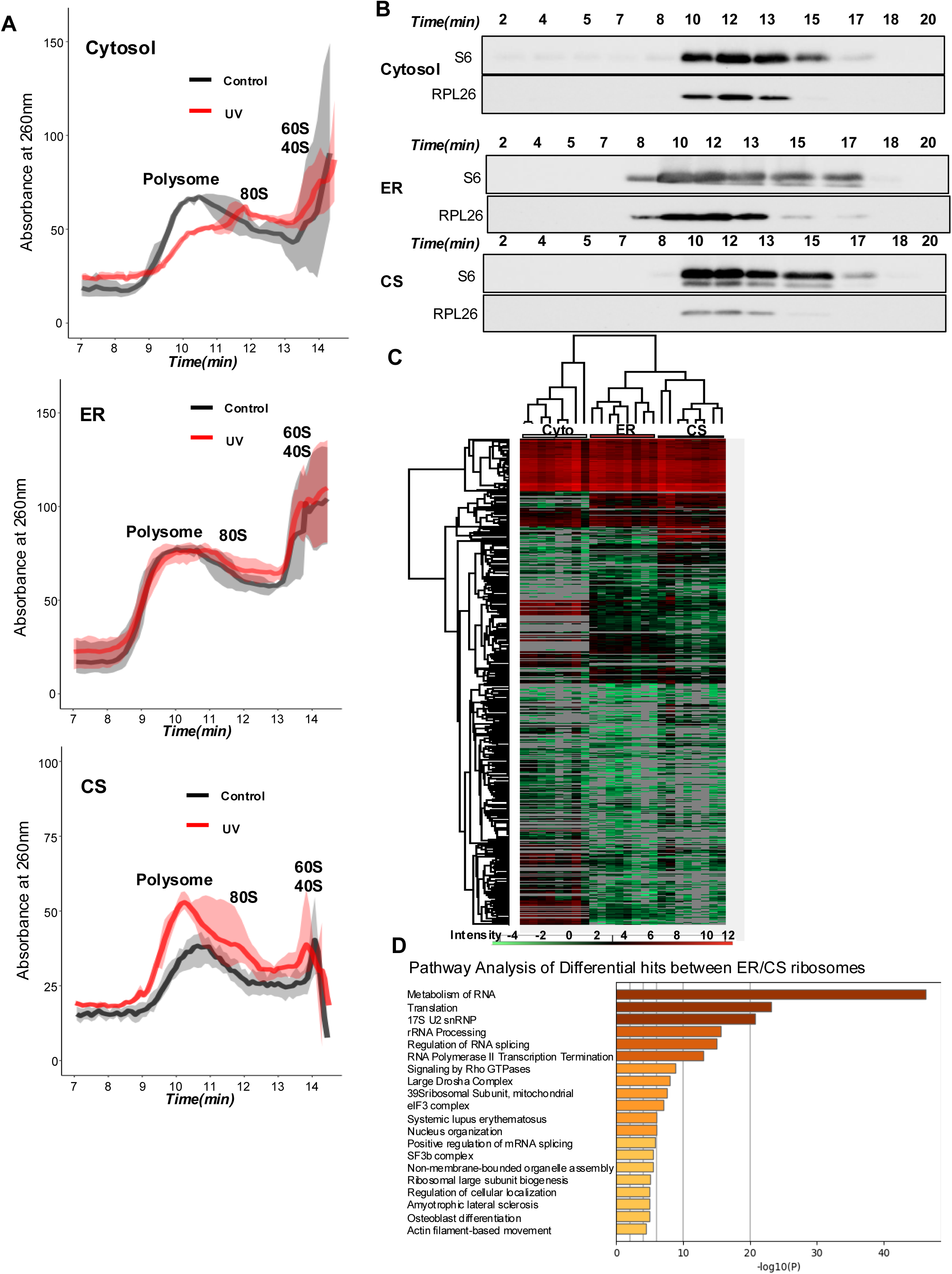
A) HPLC ribosomal profiling (RIBO-SEC) from subcellular fractions in A549 cells at baseline and after UV treatment. As described in methods, cells were first treated with UV and were then fractionated into Cytosolic, ER, and CS fractions as per fractionation protocol described in methods, followed by RIBO-SEC to obtain 260nm absorbance curves corresponding to polysomes (between 8-10.5 min), 80S (11-12 min), and 40/60S with other proteins (>13 min). The control curves are shown in black and the UV curves are shown in red. The translucent area around the curve corresponds to the variability between the biological replicates (n=3). The area under the curves for polysomal peaks is quantified as shown in Fig. 4D). Samples were run on 2000Å HPLC column, which is ideal for separating polysomes but not small structures, as seen by the decreased variance for polysomes compared to smaller structures at longer time points. B) Immunoblots for S6 and RPL26 for sequential fractions taken at different time points from RIBO-SEC experiment corresponding to the polysomal peaks in Fig. S6A from the control group for Cytosolic, ER, and CS fractions. C) Heatmap of proteome and interactome from proteomic mass spectrometry analysis of isolated ribosomes in Cytosolic, ER, and CS fractions. A549 cells from control and UV groups were first fractionated into Cytosol, ER, and CS fractions, then ribosomes were isolated from each fraction. Mass spectrometry was then used to analyze the proteomics of each ribosomal pools. Perseus was used to analyze the aligned and quantified data and generate the figures. Hierarchical clustering was then done for each ribosome sample in cytosol, ER, and CS ribosomes. n=8. Scale is shown below, grey lines indicate undetected hits. D) Pathway analysis, using Metascape, of the significant protein hits between ER and CS ribosomal pools, the two closest ribosomal clusters (Fig. S6C) using ribosome isolation and mass spectrometry. Pathway analysis reveals important differences in RNA binding proteins, translation regulation, and ribosomal structural proteins, among others, between ER and CS ribosomal proteins and ribo-interactome (i.e., proteins associated with each ribosomal pool, including binding proteins and translated proteins). Analysis is done on total significant hits.

**Figure S7.**
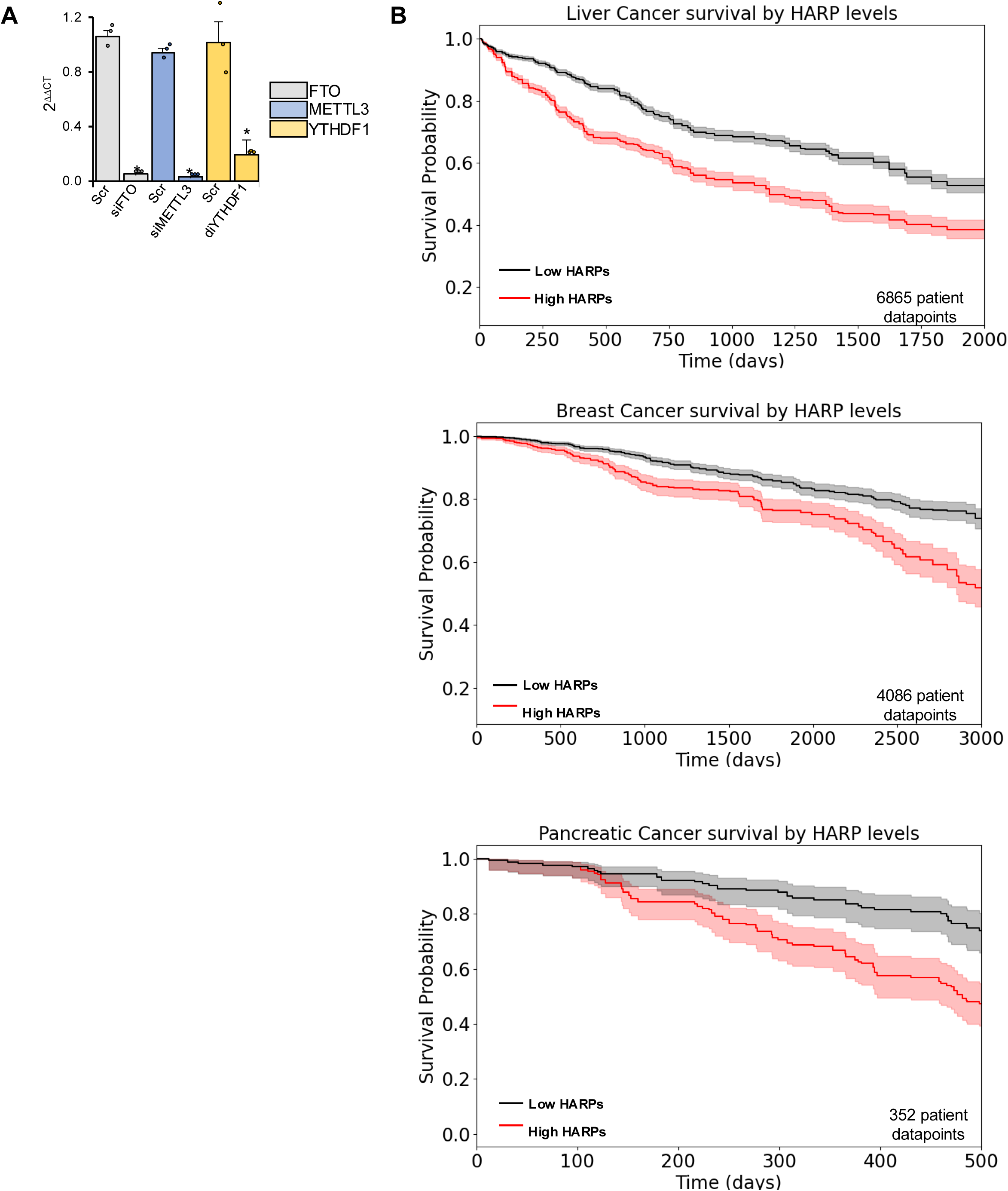
A) Mean data of mRNA levels for FTO, METTL3, and YTHDF1, measured in the siRNA treated groups (from Fig S6E) using qRT-PCR, showing decreased mRNA levels with the respective siRNA treatment. B) Kaplan Meier curves of survival probability of Liver, Breast, and Pancreatic cancer patients based on level on HIGH vs LOW HARP expression show decreased survival in HIGH compared to LOW HARP-expression group. This suggests that High levels of HARPs offer tumors more resistance to stress.

## Notes

### Competing Interest Statement

The authors have declared no competing interest.

